# The structure of the human LACTB filament reveals the mechanisms of assembly and membrane binding

**DOI:** 10.1101/2022.04.21.489104

**Authors:** Jeremy A. Bennett, Lottie R. Steward, Johannes Rudolph, Adam P. Voss, Halil Aydin

## Abstract

Mitochondria are complex organelles that play a central role in metabolism. Dynamic membrane-associated processes regulate mitochondrial morphology and bioenergetics in response to cellular demand. In tumor cells, metabolic reprogramming requires active mitochondrial metabolism for providing key metabolites and building blocks for tumor growth and rapid proliferation. To counter this, the mitochondrial serine beta-lactamase-like protein (LACTB) alters mitochondrial lipid metabolism and potently inhibits the proliferation of a variety of tumor cells. Mammalian LACTB is localized in the mitochondrial intermembrane space, where it assembles into filaments to regulate the efficiency of essential metabolic processes. However, the structural basis of LACTB polymerization and regulation remains incompletely understood. Here, we describe how human LACTB self-assembles into micron-scale filaments that increase their catalytic activity. The electron cryo-microscopy (cryoEM) structure defines the mechanism of assembly and reveals how highly ordered filament bundles stabilize the active state of the enzyme. We identify and characterize residues that are located at the filament-forming interface, and further show that mutations that disrupt filamentation reduce enzyme activity. Furthermore, our results provide evidence that LACTB filaments can bind lipid membranes. These data reveal the detailed molecular organization and polymerization-based regulation of human LACTB and provide new insights into the mechanism of mitochondrial membrane organization that modulates lipid metabolism.

## Introduction

Mitochondria are essential components of eukaryotic cells that communicate cellular needs to ensure optimal cellular function [1]. Although mitochondria play a critical role in energy production, mitochondrial metabolism is multifaceted and supports a variety of cellular functions, including free radical production, ion homeostasis, and biosynthesis of precursors for macromolecules [2]. Unlike most organelles, mitochondria are composed of two membranes: an outer membrane (OM) that interfaces directly with the cytosol, and a morphologically complex inner membrane (IM). The inner membrane separates the intermembrane space (IMS) from the matrix and folds inwards to form cristae [3,4]. This complex double-membrane architecture provides discrete compartments for mitochondria to perform diverse metabolic functions [3]. In addition to their central role in various biochemical pathways, mitochondria are remarkably dynamic organelles that constantly adopt a range of morphologies and are actively transported to specific subcellular locations in response to cellular stressors [5,6].

Mitochondria are a major site for lipid metabolism, where biosynthesis of the phospholipids such as phosphatidylethanolamine (PE), phosphatidylglycerol (PG), and cardiolipin (CL) as well as redox-active lipid coenzyme Q (CoQ, ubiquinone) is facilitated by mitochondrial proteins and protein complexes [7–9]. Mitochondrial lipids are involved in maintaining mitochondrial morphology, cristae development, organelle dynamics, regulation of membrane-associated proteins, mitophagy, and regulated cell death [10,11]. The key phospholipid PE is synthesized *de novo* by phosphatidylserine decarboxylase (PISD), a mitochondrial IM enzyme that converts phosphatidylserine (PS) to PE [11]. The newly synthesized PE is then exported to the endoplasmic reticulum (ER) for further conversion to phosphatidylcholine (PC), the most abundant membrane lipid in eukaryotic cells [11]. Disruption of the orthologous gene *Pisd* in mouse models results in embryonic lethality associated with mitochondrial dysfunction and aberrant mitochondrial morphology [12–14]. Hence, *de novo* synthesis of PE is essential for the efficiency of many subcellular processes and compartmentalization [11].

In many cancer cells, altered lipid metabolism is among the most prominent metabolic alterations that promote rapid cancer cell growth and tumor formation [15]. Processes that regulate lipid levels are important targets for the differentiation and proliferation of cancer cells [15]. Being at the junction of various metabolic pathways, mitochondrial processes are also associated with tumorigenesis and tumor progression [16]. Although mitochondrial dysfunction and aerobic glycolysis have been widely accepted as hallmarks of cancer, functional mitochondria are also important for tumor growth as active mitochondrial lipid metabolism supports the metabolic needs of highly proliferative cancer cells throughout tumorigenesis [16,17]. Conversely, tumor suppressor proteins can influence the production of lipids by acting through their biosynthesis pathway [18–20]. In mitochondria, the serine beta-lactamase-like protein (LACTB), which is localized in mitochondrial IMS, functions as a tumor suppressor by regulating the PE biosynthetic pathway [20–22]. LACTB is widely expressed in different mammalian tissues, most notably in the skeletal muscle, heart, and liver, and was found to negatively affect the proliferation of a variety of tumor cells *in vivo*, while it exerts little or no effect on the growth of non-tumorigenic cells [20,23]. It has been reported that LACTB expression has been significantly downregulated by promoter methylation, histone deacetylation, and several microRNAs (miRNAs) in various cancers, including breast cancers, uterine cancer, colorectal cancers, gliomas, melanomas, hepatocellular carcinomas, and oxaliplatin-resistant gastric cancer [24–29]. The downregulation of LACTB expression is often correlated with a poor prognosis in these anomalies [28]. While the direct mechanism for tumor suppression by LACTB is unknown, studies have shown that LACTB expression results in reduced PISD protein levels in human breast cancer cells [20]. Downregulation of PISD by LACTB is associated with markedly reduced production of PE from PS in carcinoma cells and a decrease in cell proliferation [20]. A possible mechanism for the regulation of PISD by LACTB may involve the formation of micron-scale filaments that control the distance between the inner and outer mitochondrial membranes such that the reaction to form PE by PISD1 is unable to occur due to spatial rearrangement [30]. The downregulation of LACTB mRNA and protein levels by cellular factors would result in reduced protein quantities in mitochondria, hinder filament formation, limit the length of self-assembled filaments, and decrease the catalytic activity of the enzyme *in vivo*; therefore, debilitating its regulatory role in lipid metabolism. Since PE plays an important role in promoting mitochondrial dynamics [14,31,32], the dysregulation of PE homeostasis due to a disruption to the LACTB self-assembly mechanism would be detrimental to mitochondrial function and spatial organization. However, the exact mechanism underlying LACTB on tumor suppression, the association between the downregulation of LACTB gene expression by miRNAs, and the effect on filament formation have yet to be determined. Moreover, mammalian LACTB regulates mitochondrial respiratory complex I activity [33] and participates in cellular metabolic processes [34–36]. Dysregulation of LACTB is associated with obesity and atherosclerosis potentially due to its involvement in metabolic pathways, in particular the phospholipid metabolism in mitochondria [20,34,35]. Given its strong relation to metabolic pathways, *LACTB* is also validated as an obesity gene capable of modifying adiposity [34].

LACTB is a conserved mammalian active-site serine protease present in all vertebrates [37]. The conserved catalytic serine residue within the ^164^SXXK^167^ motif plays an important role in the enzymatic activity of LACTB and is reversibly acylated through substrate binding [21,37]. Amino acid sequence analyses show that serine protease LACTB shares sequence similarity to the penicillin-binding protein and β-lactamases (PBP-βLs) family involved in peptidoglycan synthesis [37]. Given that the sequences of β-lactamases are conserved from bacteria to humans, it has been predicted that the molecular mechanisms underlying the cellular function are conserved for β-lactamases across species [37]. However, very little is known about the structural properties and function of LACTB in eukaryotic organisms. In contrast to its prokaryotic homologs, the human LACTB is expressed as a single polypeptide with an N-terminal mitochondrial targeting sequence [22,23]. Furthermore, mammalian LACTB dynamically assembles into filamentous polymers extending more than a hundred nanometers [21]. Filamentation has been observed for a variety of metabolic enzymes and is recognized as a general mechanism for regulating activity [38–45]. For LACTB, the mechanisms by which filament formation modulates enzyme activity remain unclear. Filaments of mammalian LACTB are reported to have a possible structural function in the spatial organization of mitochondrial IMS [21]. Recently, endurance exercise training has been shown to affect the polymerization of LACTB molecules in mice mitochondria suggesting that polymer assembly may have been directly influenced by metabolic activity in cells [36]. The formation of filaments may serve a regulatory function that results in more rapid activation or inhibition of the enzyme in response to changing cellular conditions.

A structure of a LACTB filament can help elucidate the biochemical characteristics of LACTB and permit a comprehensive mapping of polymerization activity. Here, we describe the atomic-resolution structure of the LACTB filament determined by using cryoEM. The structure reveals the inter-subunit interactions and overall assembly mechanism of LACTB. Furthermore, we provide supporting biochemical analysis of the assembly, where structure-guided mutations at different filament-forming interfaces unveil that polymerization is essential for the catalytic activity of LACTB. Importantly, our work reveals that LACTB exerts its regulatory role by directly interacting with lipid membranes, building a basis for understanding the mechanism of mitochondrial membrane organization. These insights expand our understanding of human LACTB function and lay the foundation for further studies of its dynamic regulation and role in tumor suppression and metabolism.

## Results

### Functional characterization of human LACTB filaments

To examine the molecular architecture of human LACTB, we purified near full-length LACTB (residues 97-547) lacking the N-terminal targeting sequence (Fig 1A-C). The protein was fractionated further with Superose 6 Increase size-exclusion chromatography (SEC), wherein a single peak corresponding to a large molecule was observed on the elution profile indicating the formation of high-molecular-weight species (Fig 1D). These oligomers seemed to be dynamic, as illustrated by the asymmetric and tailing peak on the gel filtration column (Fig 1D). Analysis of the chromatogram peak using negative-stain transmission electron microscopy (TEM) revealed that LACTB assembled into filaments of several hundred nanometers in size (Fig 1E). Our initial observations suggested that the polymers consisted of repeating LACTB subunits, resulting in flexible filaments with an outer radius of 150 Å and a wide range of length distribution (Fig 1E). To test the proteolytic activity of LACTB filaments, we performed an *in vitro* substrate assay with the fluorogenic Ac-YVAD-AMC peptide and readily detected the enzymatic activity of purified wild-type (WT) LACTB in the polymerized state (Fig 1F). These results confirm that, like the other metabolic enzymes that self-assemble into filaments to organize and regulate metabolism [38,42,43], LACTB subunits can also form catalytically active filaments in solution.

**Fig 1.**
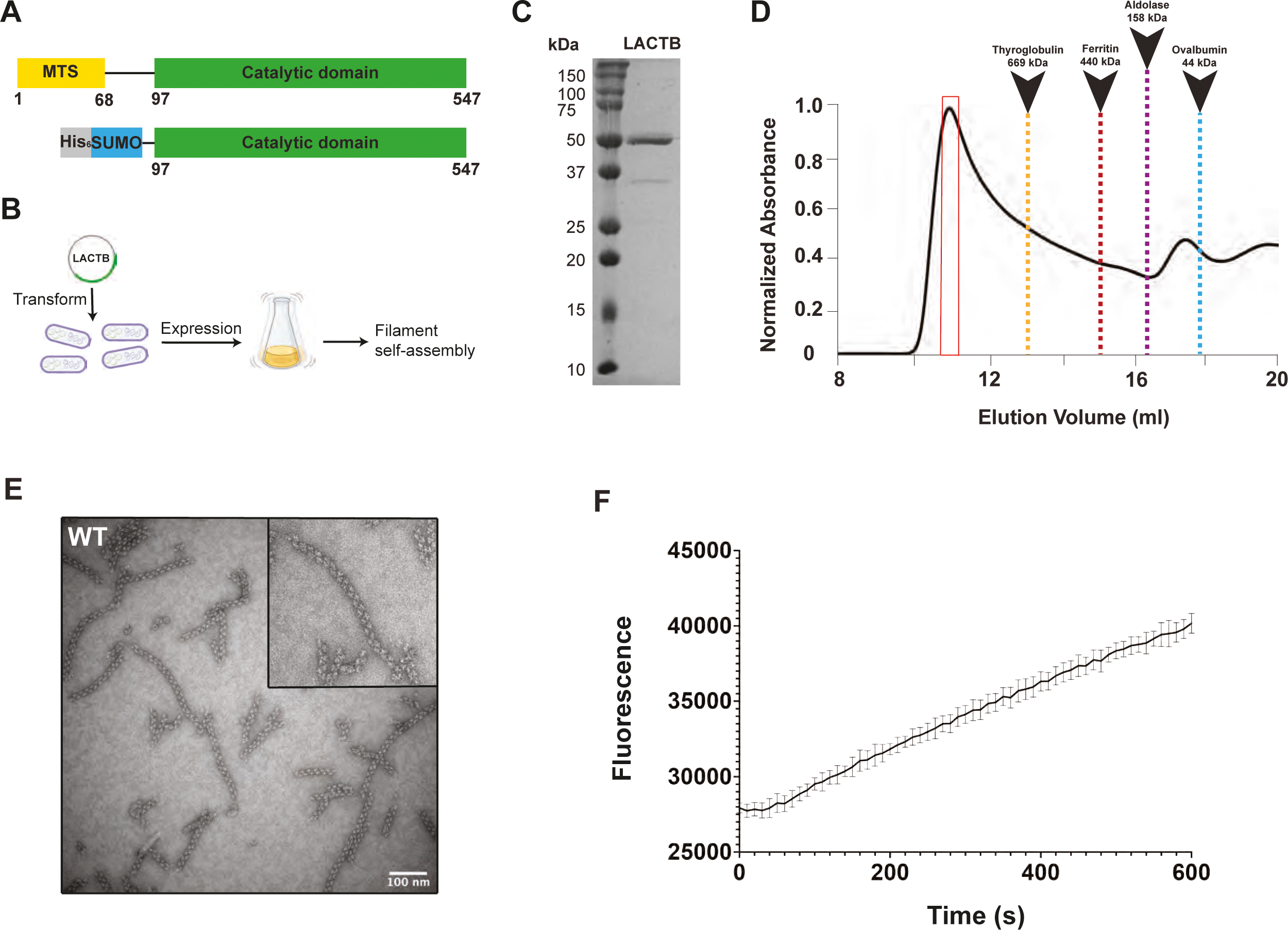
Human LACTB assembles into a filament. (**A**) Domain organization of full-length human LACTB and the construct used in the study. The color scheme is used throughout the manuscript. (**B**) Schematic diagram for recombinant expression and self-assembly of LACTB. (**C**) Coomassie-stained SDS-PAGE of purified LACTB filament. (**D**) Size-exclusion profile of WT LACTB using a Superose 6 column. SEC elution profile begins with the void volume (V_0_) of the column. (**E**) Representative negative-stain TEM image of purified samples containing high molecular weight species. Negative-stain TEM analysis is shown for the samples taken from the primary peak observed in the size-exclusion chromatogram and is highlighted with a red box in **C**. Scale bar, 100 nm. (**F**) Activity of the purified LACTB filaments was determined by using a fluorescence-based assay with the error bars representing the standard error of the mean (s.e.m.). Source data for (**F**) is provided in sheet Fig 1F in S1 Data.

### LACTB assembles into a helical filament

To establish the structural basis for the LACTB filament assembly, we used cryoEM analysis. Initial cryoEM micrographs presented substantial challenges with respect to filament flexibility and clustering under cryogenic conditions. Sample preparation was subsequently optimized by adding detergent for cryoEM grid preparation. The two-dimensional (2D) classifications of overlapping filament segments and initial three-dimensional (3D) reconstructions encompassing multiple protomers revealed the helical architecture of the filament (S1 Fig). After excluding low-quality segments, focus classification and refinement of the central region was carried out with 58% of segments, which reduced heterogeneity and improved map quality (S2 Fig). A final cryoEM reconstruction of the LACTB filament was obtained at an overall resolution of 3.1 Å with an imposed helical symmetry (S2 Fig and Table 1). A map with no noticeable differences (3.1 Å) was obtained when the final subset of segments was refined without imposing helical symmetry indicating that the helical assembly is highly symmetric (S3 Fig). The resulting map quality was sufficient for unambiguous assignment of individual subunits, and to build an atomic model of the complete structure of the LACTB filament (Fig 2 and Table 1). Except for the flexible loop region of the LACTB (residues 243 to 290), which extends to the solvent, all secondary structural elements and most side chain densities were clearly resolved in the cryoEM map (Fig 2). The flexible loop (L6) may be weakly resolved due to conformational changes between protomers or because it is mostly disordered and thus cannot be modeled. Further analysis of the structural data revealed that the basic building blocks of LACTB are dimers, which are stacked on each other and assembled into filaments with a helical twist of 48.25° and a rise of 21.63 Å (Fig 2A).

**Table 1.** CryoEM data collection, refinement, and validation statistics.

|  | Human LACTB<br>(EMD-26595)<br>(PDB 7ULW) |
| --- | --- |
| Data collection and processing |  |
| Microscope | FEI Titan Krios |
| Camera | Gatan K3 Summit |
| Magnification | 29,000x |
| Voltage (kV) | 300 |
| Electron exposure (e<sup>-</sup>/Å<sup>2</sup>) | 65 |
| Defocus (μm) | -0.5 to -1.5 |
| Pixel size (Å) | 0.8211 |
| Symmetry imposed | Helical |
| Micrographs (no.) | 7643 |
| Initial segment images (no.) | 430,592 |
| Final segment images (no.) | 248,548 |
| Map resolution (Å) | 3.09 |
| FSC threshold | 0.143 |
| Map resolution range (Å) | 3.1 to 5.8 |
| Refinement |  |
| Initial model used | *de novo* |
| Model resolution (Å) | 2.84 |
| FSC threshold | 0.143 |
| Map sharpening *B* factor (Å<sup>2</sup>) | -87.6 |
| Model composition |  |
| Nonhydrogen atoms | 18766 |
| Protein residues | 2400 |
| B factors (Å<sup>2</sup>) - min |  |
| Protein | 78.60 |
| R.m.s. deviations |  |
| Bond lengths (Å) | 0.002 |
| Bond angles (Å) | 0.406 |
| Validation |  |
| MolProbity score | 1.07 |
| Clashscore | 2.84 |
| Poor rotamers (%) | 0.00 |
| Ramachandran plot |  |
| Favored (%) | 98.74 |
| Allowed (%) | 1.26 |
| Disallowed (%) | 0.00 |

**Fig 2.**
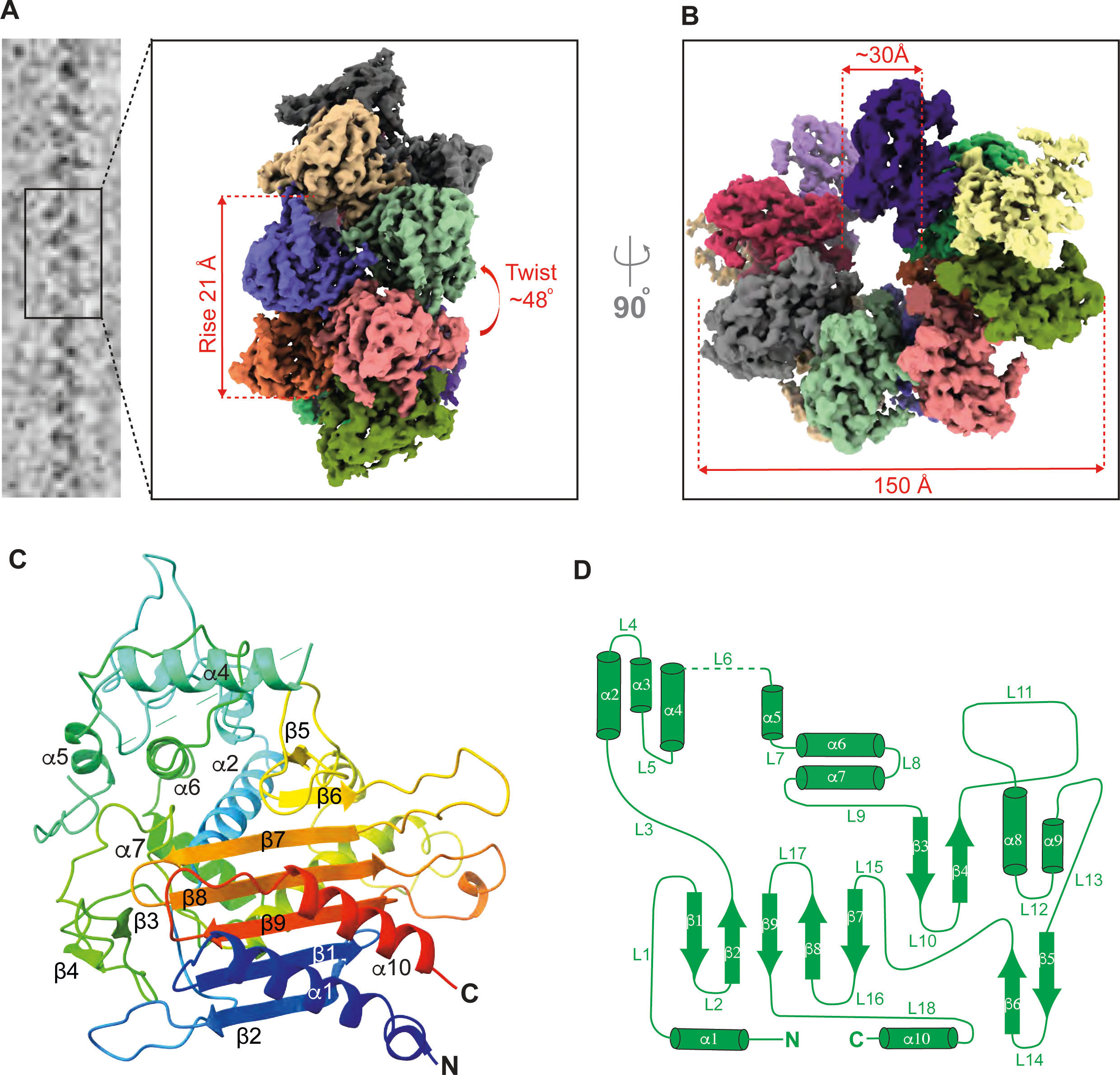
CryoEM structure of human LACTB filament. (**A**) CryoEM images of helical LACTB filaments were processed to obtain the 3D reconstruction of the polymer. LACTB protomers are colored differently, and filament rise and twist are indicated. (**B**) Top view of the LACTB assembly with the indicated filament width and inner lumen diameter. (**C**) Ribbon diagram of LACTB monomer. The chain is colored in a rainbow gradient from N-terminus (blue) to C-terminus (red). Secondary structure elements of one protomer are labeled. (**D**) LACTB (residues 97-547) α-helices (cylinders), β-sheets (arrows), and loops (lines) are depicted in a topology diagram.

### Molecular architecture of human LACTB

LACTB has a globular structure with approximate dimensions of 60 Å X 55 Å X 45 Å and consists of a five-stranded β-sheet core (β1, β2, β7, β8, and β9) surrounded by two short two-stranded antiparallel β-sheets (β3 - β4 and β5 - β6) and ten α-helices (α1 - α10) (Fig 2C and 2D). The β-sheet core is flanked by two helices (α1 - α10 and α2 - α8) on either side, creating a compact structure that forms the catalytic core (Fig 2D). These α-helices and the β-sheet core are primarily stabilized by a network of hydrophobic interactions. While the α1 and α10 helices are connected to the β-sheet core with short loops, a 21-residue loop (L3) spans across the β-sheet core to link β2 to α2, and another long loop (L11) resides between the β4 strand and the α8 helix (Fig 2C and 2D). The α-helices, α3-α7 and α9, do not directly interact with the β-sheet core and form the outer surface of the filamentous assembly (Fig 2C). Many charge-charge interactions stabilize the α-helices and the long loops of LACTB. Consistent with these observations, a Dali server search identified several prokaryotic and archaeal PBP-βL family of enzymes, such as ClbP (yellow, PDB ID: 4GDN), Pab87 peptidase (orange, PDB ID: 2QMI), Penicillin-binding protein (pink, PDB ID: 3TG9), and D-amino-acid amidase (purple, PDB ID: 2EFU), as close structural homologs of LACTB (S4 Fig). While the resemblance in the overall fold of LACTB to prokaryotic homologs suggests the conservation of the enzymatic core, significant differences are observed in the loops and α-helices that surround the core structure (S4 Fig). Most importantly, the LACTB monomer includes a long L15 loop (residues 467 to 482) formed between β6 and β7 strands, a helix α4 (residues 227 to 242) that extends in parallel with the β-sheet core, and the predicted flexible loop (residues 243 to 290), all of which are not observed in other PBP-βLs proteins (S4 Fig). Our 3.1 Å map also enabled the identification of the amino acid motifs (^164^SISK^167^, ^323^YST^325^, and ^485^HTG^487^) that contribute to the formation of the catalytic site, including the catalytic S164 residue, all of which are conserved across metazoan species (S5 Fig). The structure highlights a clearly defined cavity, which has a predominantly positively charged character (S5B Fig). The LACTB binding pocket is a ∼20 Å deep cavity with a wider outer diameter (∼20 Å) and a narrower inner diameter (∼10 Å) near the catalytic site (S5A and S5B Fig). In addition to the conserved active site residues, several positively charged (H216, H222, and K394) and aromatic residues (Y223, W454, and Y460) are in close proximity to the catalytic site (S5 Fig). This suggests that LACTB substrates largely dock through charge-charge interactions, with a few polar residues responsible for the catalytic activity.

### LACTB dimer interface governs filament assembly

We next sought to establish the mechanism of filament assembly. The LACTB filaments assemble as stacked antiparallel dimers, where the buried surface area between the two protomers (A and B) is ∼945 Å^2^ (Fig 3A). The structure shows that the antiparallel arrangement between two LACTB monomers results in symmetric interactions between two main dimer interfaces located on the L3 and L9 loops (Fig 3A). Assembly of the LACTB dimer is mediated by the combined effects of sixteen hydrogen bond interactions between E149, N150, N364, E365, P366, I368, N370, R371, N385, and T386 residues, and two inter-molecular salt bridges between E149 and R371 (Fig 3B). Conservation of these residues across mammalian species suggests that they may play an important role in subunit interactions (S5C Fig). Consistently, mutations targeting the dimer interface, including E149R-N150A double mutant and I368A-N370A-R371E triple mutant completely abolished the dimerization propensity as well as the filament formation of LACTB and produced a peak appearing at a lower molecular weight corresponding to a monomer in SEC analysis (Fig 3C). Additionally, SDS-PAGE analysis and negative-stain electron micrographs of the peak fractions further confirmed our observations on the sizing column and showed that these substitutions remarkably destabilize the local complementary interactions and thus hamper the filament formation (Fig 3C). On the other hand, mutation of N364A-E365A only modestly affected the overall filament stability and showed a gel filtration profile similar to the WT (Fig 3C). Because the N364A-E365A double mutant did not completely hinder filament formation, we predicted that the dimer interface is likely stabilized by multiple polar and electrostatic interactions. To understand how filament assembly and disassembly influence the catalytic activity of LACTB, we characterized the activity of dimerization mutants in substrate assays. Compared to the WT LACTB, the dimer interface mutants, E149R-N150A and I368A-N370A-R371E, were completely inactive, while the N364A-E365A mutation only led to a 20% reduction in LACTB activity (Fig 3D). The negative effect of every dimer interface mutation disrupting the filament assembly suggests that dimerization and subsequent polymerization are necessary to achieve maximal catalytic efficiency. These data provide a mechanism by which the strong interactions between the opposing dimer interface loops promote filament growth and result in increased enzyme activity by stabilizing the active architecture of the catalytic site. Taken together, we can confirm that the LACTB homodimers are the basic building blocks of the filaments, and polymerization is strongly correlated with optimal enzyme activity.

**Fig 3.**
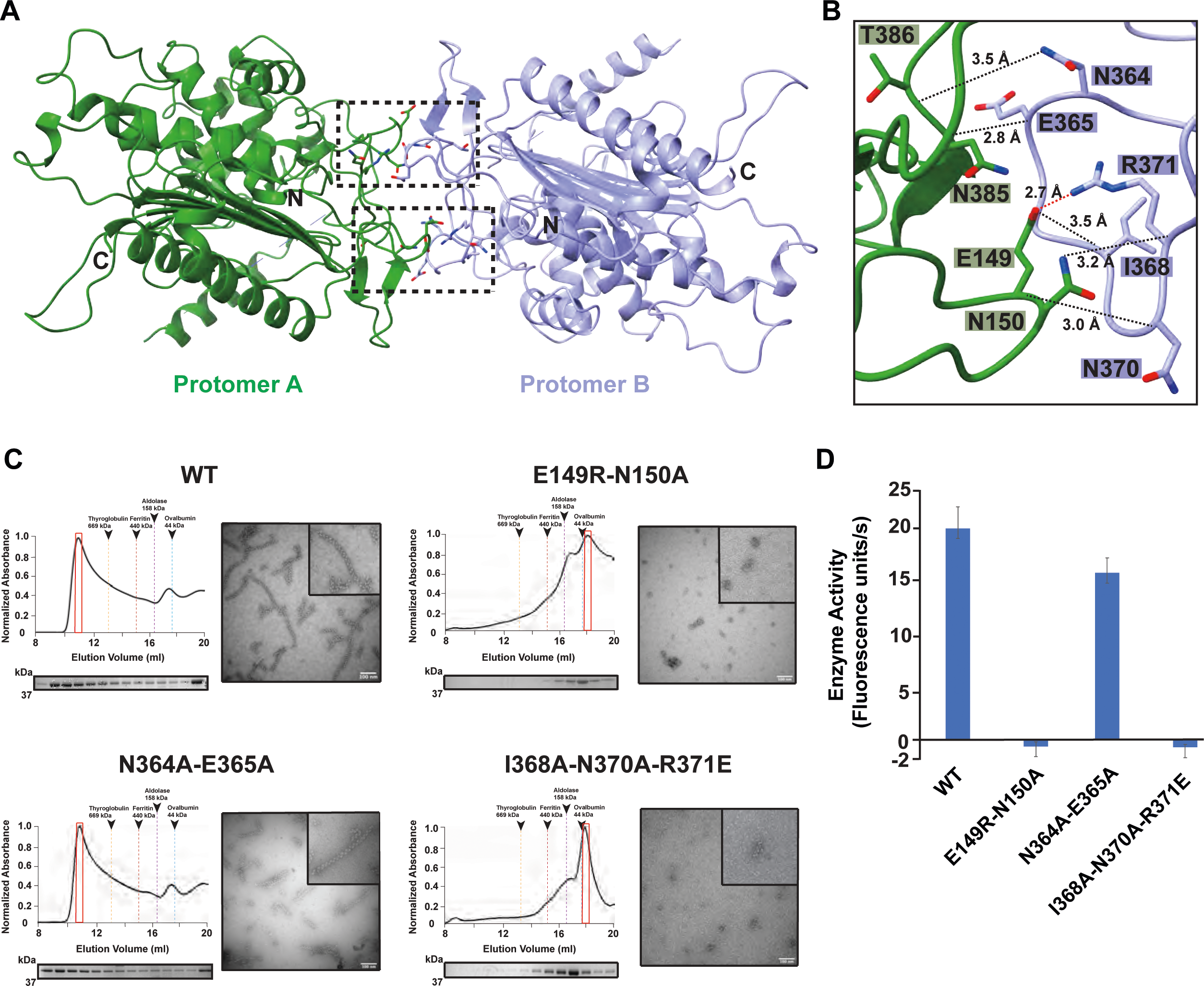
Structural characterization of LACTB dimerization interface. (**A**) LACTB protomers (colored in green and lilac) form an antiparallel dimer as shown in ribbon representation. Residues that are critical for oligomerization are represented in sticks. Black boxes indicate the position of dimer interface residues. (**B**) Zoomed in view of the black box in **A**, showing the interactions that stabilize the dimer formation. The polar and charge-charge interactions are indicated by black and red dotted lines, respectively. (**C**) Size-exclusion chromatography (SEC) elution profiles of WT and mutant LACTB. SEC elution profiles begin with the void volume (V_0_) of the column. Fractions were analyzed by SDS-PAGE and peak fractions used in negative-stain TEM analyses are highlighted with red boxes. Negative-stain TEM images demonstrate the polymerization activity of WT and mutant proteins. Scale bars, 100 nm. (**D**) Mutation of key residues involved in dimerization disrupts catalytic activity. Enzyme activity was determined *in vitro* by using a fluorescently labeled substrate and three independent experiments were performed for each sample. Bars represent the mean of at least three independent experiments and error bars represent the standard deviation. The underlying data in (**D**) is provided in sheet Fig 3D in S1 Data.

### LACTB filament elongation is mediated by multiple polymerization interfaces between subunits

When we evaluated multiple LACTB homologs, we decided to focus our structural analysis on the unique polymerization mechanism that brings the LACTB homodimers together to form the full filament. Polymerization is mediated by the association between two LACTB homodimers (protomers A-B and C-D), where 48.25° helical rotation between stacked homodimers is necessary for growing a filament at either end (Fig 4A). The inspection of the inter-dimer interface shows that three assembly contacts between neighboring subunits (A and C, A and D, and B and D) bridge antiparallel LACTB homodimers and drive filament assembly (Fig 4A). The first polymer interface, which spans a buried solvent-accessible surface area of ∼780 Å^2^, is formed between the α1 helix and L3 along with the protruding L10 loop between β3 and β4 strands of protomers A and D (Fig 4A and 4B). The two-fold symmetric interactions between protomers A and D from neighboring homodimers result in identical electrostatic surface potentials on both sides of the inter-dimer interface (Fig 4A and 4B). Major contacts of this interface include several non-covalent interactions between residues D113, R117, E121, V122, R151, Y377, R382, L383, and S527, and a pair of symmetrical salt bridges between D113, D120, and R151 (Fig 4A and 4B). To verify the functional relevance of these interactions for polymerization and catalytic activity, we performed mutagenesis studies on residues R117, R151, and Y377. Both R117E and R151E charge-reversal mutations hindered the ability of LACTB to assemble into stable elongated filaments and yielded shorter assemblies containing a few LACTB protomers (Fig 4D). Notably, the R151 salt bridge mutation reduced the proteolytic activity of LACTB by ∼60% in substrate assays, whereas the R117E mutation only mildly affected the enzyme activity (Fig 4E). Moreover, mutation of the conserved contact site residue Y377 to a lysine did not show major changes in filament length and formation compared to the WT but resulted in a greater effect (∼70% reduction) regulating protease activity (Fig 4D and 4E). These characteristics, together with the disruptive effect of charge-reversal mutations, indicate that complementary electrostatic forces stabilize the interactions between stacked LACTB dimers during filament formation.

**Fig 4.**
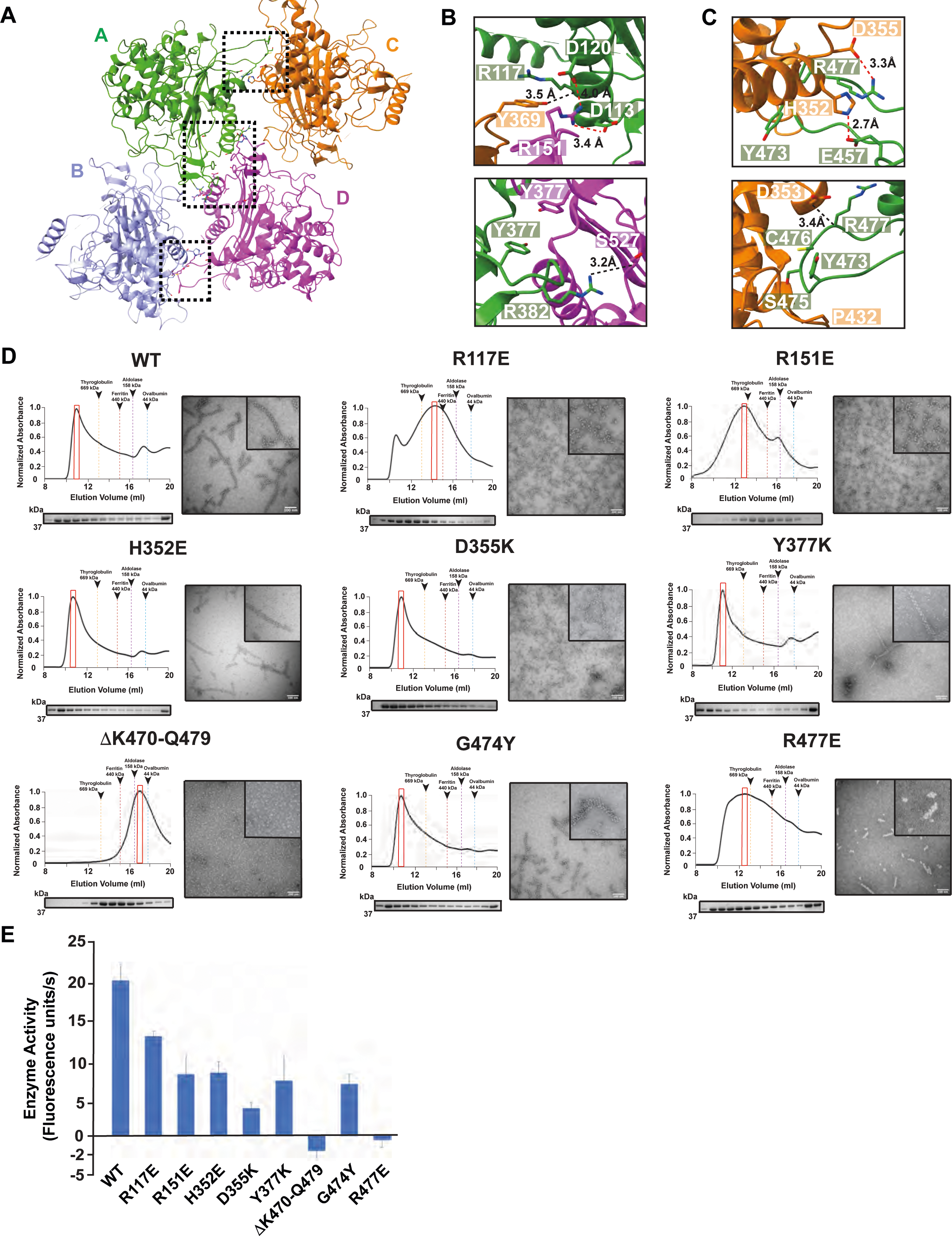
Structural basis of LACTB filament assembly. (**A**) Ribbon diagram of LACTB tetramer. LACTB protomers are colored in green (A), lilac (B), orange (C), and pink (D). Residues that are critical for oligomerization are represented in sticks. Black boxes indicate the position of the residues located at both polymerization interfaces 1 and 2, respectively. (**B** and **C**) Zoomed in view of polymer interfaces 1 and 2, respectively, showing the interactions that facilitate the polymerization of human LACTB. The polar and charge-charge interactions are indicated by black and red dotted lines, respectively. (**D**) Size-exclusion chromatography (SEC) elution profiles of WT and mutant LACTB. SEC elution profiles begin with the void volume (V_0_) of the column. Mutations to residues that are located at the polymerization interfaces 1 and 2 impair filamentation. Fractions were analyzed by SDS-PAGE and peak fractions used in negative-stain TEM analyses are highlighted with red boxes. Negative-stain TEM images demonstrate the polymerization activity of WT and mutant proteins. Scale bars, 100 nm. (**E**) Mutation of key residues involved in filament formation disrupts catalytic activity. Enzyme activity was determined *in vitro* by using a fluorescently labeled substrate and three independent experiments were performed for each sample. Bars represent the mean of at least three independent experiments and error bars represent the standard deviation. The source data from (**E**) is provided in sheet Fig 4E in S1 Data.

The filament structure also shows a second polymer interface that has symmetric contact sites between protomers A and C and B and D, which bury ∼540 Å^2^ of solvent-accessible surface area (Fig 4A). These two interfaces are formed by the same residues and the interactions are mainly mediated by the charge complementarity and salt bridges (Fig 4A-C). In our filament structure, the unique L15 loop that extends between β6 and β7 strands is embedded within the large pocket formed by α7 and α8 helices and the L12 loop of the adjacent LACTB protomer (Fig 4A-C). The most intimate contacts occur between L15 loop residues Y473, G474, S475, C476, and R477, and the conserved residues H352, D353, D355, Y369, A413, Y416, P432, and Y434 that line the surface of the pocket (Fig 4C). Additional inter-molecular salt bridges are formed between H352 and E457, and D355 and R477, which stabilize the filament formation (Fig 4C). To assess the importance of these interfaces, we produced a LACTB construct in which the L15 loop (V467 to Y482) is replaced by a glycine-glycine-serine (GGS) linker to disrupt the helical assembly interface. This mutant impaired the ability of LACTB to form long filaments, as judged using size-exclusion chromatography and negative-stain TEM (Fig 4D). We then tested this construct in substrate assays to measure catalytic activity. Not surprisingly, the removal of the L15 loop completely abolished the enzymatic activity of the purified LACTB oligomers (Fig 4E). Mutations of individual L15 loop residues also hindered the enzymatic activity of LACTB. Substitution of residue G474 with tyrosine markedly reduced the catalytic activity of LACTB by ∼85%, whereas charge-reversal mutation of R477 residue resulted in abrogation of catalytic function *in vitro* (Fig 4E). We also characterized the role of the inter-molecular charge-charge interactions at the interface. Mutations targeting interface salt bridges, including H352A and D355K, did not affect the filament formation but decreased the enzymatic activity of LACTB by ∼60% to 80%, respectively (Fig 4D and 4E). Although these interface residues do not appear to participate in catalytic site formation as they are distant from the active site, they facilitate inter-subunit interactions that increase the stability and catalytic activity of the filament. Interestingly, a structural comparison of the human and prokaryotic PBP-βL family of enzymes unveiled that the LACTB structure is distinguished from its prokaryotic homologs by the addition of the 14-residue L15 loop, which is involved in filament assembly (S4 Fig). Moreover, multiple sequence alignment of the human LACTB and PBP-βL family of enzymes demonstrated very little sequence similarity for amino acids forming the L15 loop (S4E Fig). Consistent with our functional analysis, the L15 loop emerges as a regulatory element that controls not only the polymerization but also the catalytic activity of LACTB. Altogether, these results demonstrate that assembly of the filament and the L15 loop are required for catalytic activation of LACTB.

We next tested the possibility that changes in reactive oxygen species (ROS) in the mitochondrial IMS may influence the assembly of LACTB filaments. Mitochondrial ROS regulate a wide variety of cellular processes and are linked to multiple pathologies [46,47]. It was previously shown that hydrogen peroxide (H_2_O_2_) can disarrange actin polymers to provoke cell death during inflammation [48]. H_2_O_2_ has also been linked to the regulation of mitochondrial membrane morphology and processes of fusion/fission by changing the oligomeric state or monomeric structure of redox-sensitive proteins [49–51]. In our cryoEM structure, we identified a methionine-rich region, where more than half of the methionine residues in LACTB are clustered in this region (S6A Fig). We thus postulated that these methionine residues may be sensitive to H_2_O_2_-mediated oxidation, which could disrupt filament assembly. To investigate the impact of mitochondrial ROS on LACTB, we exposed protein samples to varying concentrations of H_2_O_2_ (0.01% to 1%) and showed that LACTB samples oxidized with H_2_O_2_ migrate more slowly than the native protein as deduced by SDS-PAGE analysis (S6B Fig). Subsequently, we confirmed the oxidation of these methionine residues upon exposure to H_2_O_2_ by liquid chromatography with tandem mass spectrometry (LC-MS/MS) (S6C Fig). Finally, negative-stain TEM imaging of protein samples revealed that oxidation at higher H_2_O_2_ concentrations hinders the filament formation of LACTB (S6D Fig). These data suggest that the H_2_O_2_-mediated oxidation of the protein could alter LACTB self-assembly dynamics and inhibit polymerization *in vitro*.

### Mechanism of LACTB membrane organization

LACTB has been shown to localize to the mitochondrial IMS, where it assembles into polymers to regulate lipid metabolism [20–22]. The structure of the LACTB filament can inform its mechanism of action and interactions with lipid membranes. To investigate whether there is a link between mitochondrial membrane organization and LACTB filament formation, we reconstituted freshly purified LACTB with liposomes that closely mimic the lipid composition of mitochondrial inner membranes in a buffer of physiological ionic strength and visualized binding *in vitro*. When analyzed by negative-stain TEM, we directly observed the interplay between LACTB and lipid membranes (Fig 5). LACTB filaments exhibited well-ordered, tightly packed conformation, and appeared to interact with liposomes via the tip of the polymer (Fig 5A). These filaments were often attached to two different liposomes on opposite ends forming a bridge between the two lipid membranes (Figs 5B-E and S7D). Intriguingly, we also identified filaments that interact with liposomes from the side, where multiple protomers form intimate interactions with lipid membranes on either side (Figs 5C-D and S7D). This second binding mode allows LACTB filaments to accommodate different membrane topologies and tightly associate with surrounding membranes, which enables membrane organization. These observations complement data that were previously obtained *in situ* [21], indicating that recombinant LACTB generates biologically relevant assemblies and binds membranes *in vitro* (Fig 5). To further characterize the molecular details of protein-lipid interactions, we first measured the enzymatic activity of LACTB in the presence of CL-containing liposomes and uncovered that membrane binding does not affect the catalytic activity of LACTB filaments (Fig 5G). Subsequently, we prepared liposomes with four different lipid compositions to test the effect of mitochondrial IM lipids on LACTB activity (S7A Fig). We observed an average two-fold decrease in liposome binding when we omitted CL from lipid vesicles (S7B-C Fig). Moreover, human LACTB filaments facilitate mitochondrial membrane organization, which requires these filaments to form stable interactions with two distinct membrane regions and coordinate their spatial organization [21]. To assess this activity *in vitro*, we also calculated the percentage of filaments that appeared to be tethered to two or more liposomes and observed a four-fold decrease in the tethering activity of LACTB filaments in the absence of CL (Fig 5H). We also determined whether LACTB filaments can sense regions of high membrane curvature that may be important for membrane remodeling processes. Specifically, we prepared CL-containing cylindrical liposomes that form membrane tubules and provide curved membrane surfaces. The reconstitution assays with cylindrical nanotubes revealed that LACTB filaments have a stronger affinity to cylindrical nanotubes than spherical vesicles (S7E). Together, these results indicate that CL and positively curved membranes enhance the membrane binding activity of human LACTB.

**Fig 5.**
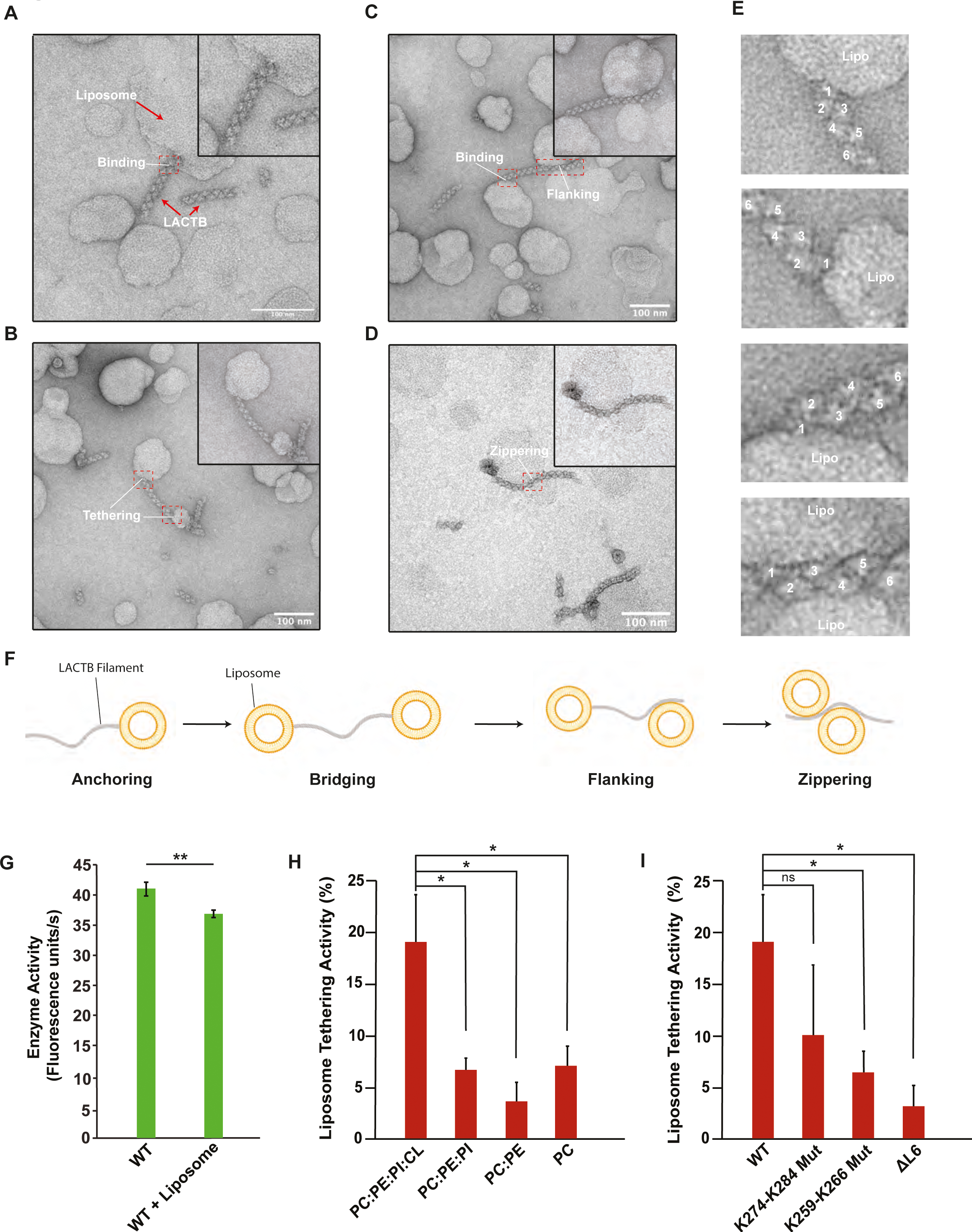
LACTB filaments bind to cardiolipin-enriched liposomes. (**A-D**) Representative negative-stain TEM images of LACTB filaments in the presence of liposomes. Scale bar, 100 nm. Purified LACTB filaments were incubated with cardiolipin-containing liposomes at room temperature for 4 hours. Red boxes indicate protein-membrane contact sites. (**E**) Zoomed in views of the red boxes in **A**, **B**, **C**, and **D**, showing the details of protein-liposome interactions. (**F**) A schematic diagram of filament-liposome interactions illustrating different membrane binding modes of LACTB filaments. (**G**) Catalytic activity of WT LACTB in the presence of liposomes. Enzyme activity was determined *in vitro* by using a fluorescently labeled substrate and three independent experiments were performed for each sample. Bars represent the mean of at least three independent WT and reconstitution assays. Error bars indicate standard deviation (**H**) Liposome tethering activity of WT LACTB under the presence of liposomes of various compositions. Liposome tethering activity was quantified by counting 1000 filaments from at least three independent reconstitution assays and calculating the total percentage of filament tethering. Bars represent the mean liposome tethering activity and error bars indicate s.e.m. (**I**) Liposome tethering activity of key residues involved in liposome binding. Liposome tethering activity was quantified by counting 1000 filaments from at least three independent reconstitution assays and calculating the total percentage of filament tethering. Bars represent the mean liposome tethering activity of reconstitution assays and error bars indicate s.e.m. Statistical analysis was performed with an unpaired two-tailed Student’s t-test (**G**) and unpaired two-tail Welch’s t-tests (**H**, **I**). *p <0.05, **p <0.005, ns (no significance). The underlying data in (**G**), (**H**), and (**I**) are provided in sheets Fig 5G, Fig 5H, and Fig 5I in S1 Data.

Furthermore, we postulated that the flexible loop region between residues 243-290 could facilitate membrane binding. Specifically, the human LACTB sequence contains a charged and hydrophobic motif (^259^KNxFxKFK^266^) as well as a cluster of six basic side chains (^274^KxRxxKxxKKK^284^) with a remarkably electropositive character within this region that may form interactions with CL-containing liposomes *in vitro* (S5C Fig). To define the molecular determinants of membrane binding, we replaced the flexible loop region (residues K255 to K284) with a glycine-glycine-serine (GGS) linker and introduced alanine and charge-reversal mutations to the ^259^KNxFxKFK^266^ and ^274^KxRxxKxxKKK^284^ motifs. These mutants were catalytically active and did not affect the ability of LACTB to form filaments, as assessed using negative-stain TEM (S8 Fig). Of note, the flexible loop deletion (ΔL6) and ^274^ExExxExxEEE^284^ mutant lowered the enzymatic activity of LACTB filaments while the ^259^EAxAxEAE^266^ mutant displayed increased catalytic activity (S8B Fig). More importantly, the ΔL6 and the ^259^EAxAxEAE^266^ mutants reduced the membrane tethering ability of LACTB filaments by more than 70% while the ^274^ExExxExxEEE^284^ mutation led to a ∼50% decrease in membrane tethering activity (Fig 5I). We also corroborated these results with computational modeling (S9 Fig). In our cryoEM map, we did not observe clearly resolved densities for the flexible L6 loop region of LACTB. Therefore, we obtained the AlphaFold predicted structure of the human LACTB monomer and calculated the rotational and translation position of the protein in the presence of membranes using the PPM (<u>P</u>ositioning of <u>P</u>roteins in <u>M</u>embranes) server (S9 Fig). These theoretical models also nominated the ^259^KNxFxKFK^266^ motif as key residues involved in membrane tethering, supporting our biochemical findings (S9 Fig). While previous studies claim that this loop may promote self-assembly of the LACTB polymer through a coiled-coil segment [21], we propose that the flexible loop on the fully solvent-exposed exterior of the filament may adopt multiple conformations to accommodate different topologies and curvatures of opposing lipid membranes, thereby mediating protein-lipid interactions that would facilitate membrane organization. Consistent with this, the flexible loop is not conserved amongst the prokaryotic and archaeal PBP-βL family of enzymes, which are not associated with lipid membranes [37], providing additional support for the relevance of this region for membrane binding (S4E Fig). In conclusion, LACTB filaments form stable interactions with cardiolipin-enriched liposomes, display different modes of membrane binding, and the key L6 loop is required for membrane targeting of LACTB filaments.

### Molecular interpretation of disease-associated mutations

Given that human LACTB exerts tumor suppression activity in multiple carcinogenic cell lines, we next investigated cancer-specific missense mutations localized within LACTB. We classified LACTB disease mutations in terms of their localization to subunit-subunit interfaces and characterized their effect on the catalytic activity (Fig 6). Currently, over 200 *LACTB* gene variants have been identified, of which ∼30% are considered pathogenic [52,53]. About 30% of the pathogenic variants cause frameshifts and premature truncation of the open reading frame, whereas the remaining ∼70% are missense mutations that are linked to various cancers [52,53]. Mapping of disease-causing mutations onto our structure of human LACTB demonstrated that some of the pathogenic variants are clustered at the oligomerization interfaces, possibly culminating in a negative effect on filament assembly (Fig 6 and S1 Table). Several mutations related to various types of cancers cluster at the LACTB dimer interface, including V148F, E149Q, E363K, A372T, R371K, K380N, and R382C variants (Fig 6B). Our biochemical characterizations showed that the E149R and R371E charge-reversal mutations resulted in a profound multimerization defect and hindered the catalytic activity of human LACTB (Fig 3C and 3D). Of note, V148, E149, and R371 belong to the L3 loop at the dimerization interface, where E149 and R371 form hydrogen bonds and an inter-molecular salt bridge to stabilize the dimer (Figs 3B and 6B). Moreover, other variants such as R151S, D457K R469K, T472K, R480L, and Y482H, which are located at the polymer filament-forming interfaces, have also been associated with clinical anomalies (Fig 6). As described earlier, we analyzed the assembly properties of the LACTB R151E mutant *in vitro* and found that this mutant depressed the multimerization of human LACTB (Fig 4D-F). Notably, we showed that the disruption of R151-D113 or H352-D457 electrostatic interactions results in a ∼60% reduction in the catalytic activity of human LACTB (Fig 4D and 4E). We observed similar results when we assessed the functional role of protruding L15 loop (residues 467 to 482), which contains several disease-associated residues that are critical for filament formation and membrane organization activity of LACTB (Fig 4D and 4E). Overall, mutations in these newly evident interfaces would likely have an impact on LACTB polymerization and catalytic activity and this provides important structural context as pathogenic variants in these interfaces are linked to several cancers.

**Fig 6.**
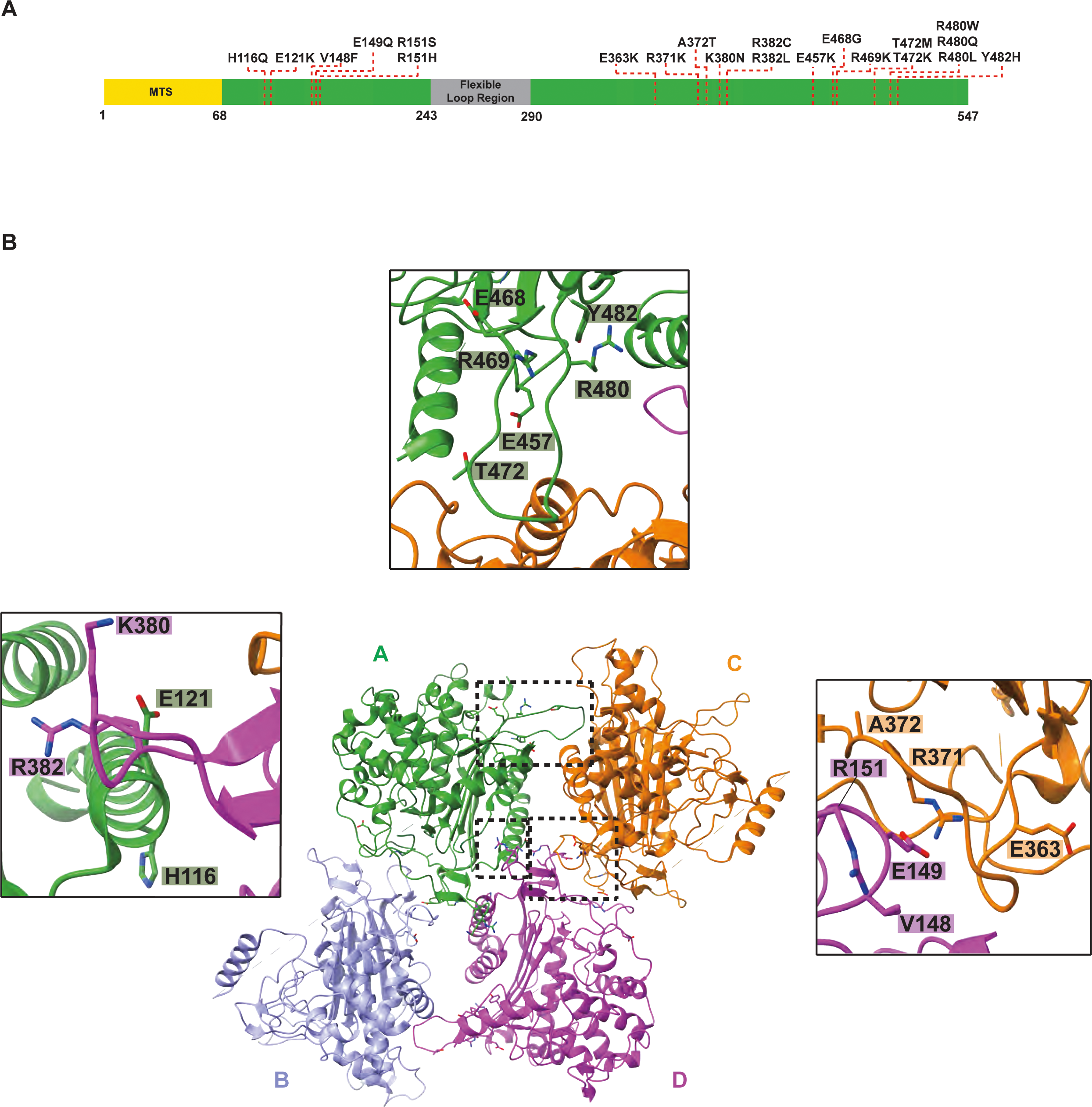
The model of LACTB enables mapping the disease-associated mutations. (**A**) A schematic representation of full-length LACTB architecture. Residues 1-68 contain the predicted mitochondrial targeting sequence (MTS), and residues 243-290 form the flexible loop region. Residues that are associated with various human cancers are highlighted with a red dashed line. (**B**) Ribbon diagram of human LACTB tetramer. Protomers that form the tetramer are labeled A to D and colored as in Fig 4A. Mutations localizing to oligomerization interfaces are illustrated as sticks. Inset windows show the close-up views of oligomerization interfaces containing disease-associated mutations. Black boxes indicate the dimer, and polymerization interfaces 1 and 2, respectively.

## Discussion

Here, we investigated the molecular mechanism of the LACTB serine protease, whose key functions in mitochondria include the regulation of metabolic pathways and morphology and tumor suppression by modulating mitochondrial lipid metabolism. Our high-resolution structure of the human LACTB filament shows that the enzyme forms a high-order helical filament, which is critical for its cellular function. These results offer new mechanistic insights into how LACTB assembles into filaments to control mitochondrial morphology as well as the efficiency of metabolic processes.

The basic building block of the helical structure consists of an antiparallel LACTB dimer. This unit then interacts with neighboring homodimers via charge-charge and polar interactions, which results in the helical stacking of LACTB dimers and elongation of filaments (S10 Fig). We show that human LACTB contains novel structural features that are critical for polymerization. Although the active site motifs are conserved between LACTB and the prokaryotic PBP-βL family of enzymes, phylogenetic and structural analyses demonstrate that the residues involved in polymerization are only conserved in higher eukaryotes, suggesting that LACTB has evolved from its prokaryotic counterparts to fulfill different physiological functions in mammals (S5 Fig). Intriguingly, LACTB requires unique contacts to robustly polymerize into filaments which enables catalytic activity. While the benefits of polymeric enzymes are not fully understood, there are several theories of filamentous enzymes exhibiting kinetic advantages. One such kinetic model involves the rapid activation of enzymes through filament formation [54–57]. For instance, the bacterial SgrIR protein, which is a restriction endonuclease, polymerizes rapidly at either end of the filament to stimulate enzyme activity and increase DNA-binding specificity [58,59]. Similarly, many metabolic enzymes such as cytidine triphosphate synthase (CTPS) and inosine monophosphate dehydrogenase (IMPDH) form filaments in bacteria and eukaryotes to modulate enzyme activity [38–41]. Interestingly, CTP-induced polymerization of bacterial CTPS results in decreased catalytic activity, whereas substrate-bound human CTPS filaments display increased enzymatic activity, suggesting that filament formation can tune the activity of metabolic enzymes differently in response to distinct metabolic cues in different species [38]. Additionally, an isoform of human acetyl-CoA carboxylase in the mitochondria forms active and inactive helical filaments through allosteric regulation by citrate and the binding of BRCA1, respectively, revealing a unique form of enzyme regulation via large scale conformational changes in filament architecture [42]. The structure of the LACTB filament, together with extensive mutagenesis studies, show that polymerization is essential for the catalytic activity of the enzyme (Fig 3). However, the polymerization interfaces do not participate in active site formation (Fig 4). A possible explanation for this increased enzymatic activity upon filament assembly is that the binding of LACTB protomers indirectly stabilizes the regions involved in substrate binding and thus enhances the catalytic activity of LACTB. The stabilization of the LACTB filament through multiple intra- and inter-dimeric interfaces that are largely mediated by charged and polar interactions strongly suggest that the filament assembly is responsible for the observed amplification of LACTB activity in our biochemical data. Our results suggest that the catalytic activity of human LACTB is allosterically regulated by stabilizing an intrinsically inactive state upon filament formation, independent of substrate binding, thus holding the enzyme in an active conformation. Filamentation appears to be an evolutionarily conserved mechanism in a diverse array of biochemical and biological pathways in cells and provides an added layer of regulation in the form of rapid activation or inhibition [56,60]. By characterizing distinct interactions in LACTB filaments our data provides a structural basis for manipulating LACTB polymerization *in vivo*.

Many different enzymes dynamically form filaments in response to changing physiological conditions in cells [38,41,42,44,61]. The process of polymerization requires a critical concentration of monomeric protein for filament formation, where larger self-assemblies can be formed as a result of protein crowding [56]. In addition to modulating enzyme activity through polymerization, an unexpected feature of the LACTB filaments is their ability to interact with lipid membranes. Our results indicate that both LACTB-mediated mitochondrial membrane organization and regulation of lipid metabolism may be directly linked to its ability to form higher-order assemblies. However, it is not clear whether the catalytic activity of LACTB plays a role in membrane organization in mitochondria as its direct substrate remains unknown. Unlike many membrane-associated proteins such as conserved septin proteins in mammals [62], membrane binding does not influence LACTB polymerization through the end-to-end association of the homodimers. It is striking that while many regulatory enzymes polymerize to allosterically tune enzyme kinetics [56], LACTB exerts a dual function and forms exclusively cooperative filaments to interact with mitochondrial membranes from the inside. We showed that the LACTB filaments are unique as they contain specific sequences to interact directly with membranes (Fig 5). The membrane topologies observed in electron tomograms of mitochondria suggest that LACTB filaments, which have the ability to access length scales larger than the size of mitochondrial membrane compartments, can form structural and functional bridges between mitochondrial IM and OM and regulate the distance between the opposing faces of the cristae membrane [63]. Hence, LACTB assemblies may have further roles in mitochondrial morphogenesis and in processes, where strong membrane association is needed. Mitochondrial membranes have a unique lipid composition that is important for the organelle’s morphology and proper function of membrane-associated processes [7]. In addition to the major phospholipids PE and PC, which account for a large portion of mitochondrial membrane lipids, mitochondria-exclusive phospholipid cardiolipin (CL) is a distinguishing component of the mitochondrial IM that is known to participate in a wide range of mitochondrial processes [10]. CL directly interacts with positively charged surfaces of numerous mitochondrial proteins with its negatively charged headgroup and contributes to the dynamic membrane organization in mitochondria [64,65]. Negative-stain TEM imaging revealed that LACTB filaments have a high affinity for cardiolipin-containing lipid bilayers and manifest two different membrane binding modes (Fig 5). We postulate that LACTB filaments are recruited to the inner/outer membrane by electrostatic interactions between negatively charged phospholipids and the flexible polybasic surface loop of LACTB. When the tip of the LACTB filaments binds lipid membranes, subunits of adjacent homodimers form additional protein-lipid interactions through lateral associations, increasing the filament contacts with the membrane surface. Furthermore, subunits on the opposite end of the filament bind membranes in a similar fashion and engage in lateral associations via interactions of several subunits, thus bringing the two membranes into close apposition. We, therefore, propose that upon forming filaments between the two membranes, human LACTB can act as a molecular zipper and regulate mitochondrial membrane organization by controlling the distance between the two membranes. LACTB filaments display a strong driving force for the spatial arrangement of membranes, thus providing a structural foundation for a wide variety of normal and pathological processes.

Finally, our ability to map mutations that are identified in various cancers to the LACTB structure expands the breadth of deficiency mechanisms (beyond the downregulation of LACTB by several miRNAs) to additionally include the impaired ability of LACTB to correctly assemble into a filament. Our structural model encompasses nearly all human cancer-associated missense mutations in LACTB and lays the foundation for characterizing additional mechanisms leading to LACTB perturbation in the regulation of lipid metabolism and mitochondrial membrane organization. This study uncovers an important aspect of how LACTB functions as a tumor suppressor and provides a foundation for future cancer research and drug therapy with LACTB as a target.

While this manuscript was under preparation, Zhang *et al.* determined the single-particle cryoEM structure of the LACTB and studied the catalytic activity of the enzyme [66]. Using topology-independent structure comparison, we determined 98.74% alignment and a root mean square deviation (r.m.s.d.) of 0.48 Å over 389 Cα atoms between the two monomeric structures. Although the overall structure of the monomeric enzyme is similar, Zhang *et al.* studied smaller LACTB oligomers that do not form elongated filaments. Our cryoEM structure of LACTB filament mirrors data that were previously obtained *in situ* [21], emphasizing that structure determination of self-assembled micron-scale filaments generates biologically relevant atomic models. Overall, our structural models and biochemical characterizations of assembly interfaces largely agree with the recently published study [66]. Moreover, our results on the assembly mechanism of LACTB and several other structural and functional analyses, including the details of protein-lipid interactions, which are critical for characterizing the regulatory role of a filamentous enzyme, provide an important basis for the elucidation of the mechanisms of LACTB protease on a molecular level.

## Materials and Methods

### Protein expression and purification

The gene encoding human LACTB (residues 1-547) was obtained from DNASU (HsCD00629745). The DNA sequence for human LACTB (residues 97-547) was subcloned into the multiple cloning site of the pCA528 bacterial expression vector. LACTB was tagged with an N-terminal 6x-His tag followed by a SUMO tag. All mutant constructs were generated by a two-step site-directed mutagenesis PCR protocol. The plasmid was transformed into *Escherichia coli* BL21(DE3)-RIPL cells and expressed in ZYP-5052 auto-induction media at 18°C for 16 h when OD_600_ reached 0.6-0.8. Cells were harvested with centrifugation (Sorvall RC-5B Superspeed Centrifuge) at 10,800 rpm (39,531x *g*) for 20 min at 4 °C. The supernatant was then discarded, and the cell pellets were resuspended in lysis buffer (50 mM HEPES-NaOH, pH 7.5, 500 mM NaCl, 5 mM MgCl_2_, 20 mM imidazole, 1mM CHAPS (Anatrace), 5mM 2-mercaptoethanol, 10% (v/v) glycerol) supplemented with 0.5% (v/v) Triton X-100, 0.5 mg DNase I, and 1X EDTA-free complete protease inhibitor cocktail (Roche). The cells were lysed with an Emusliflex C3 Homogenizer and the cell lysate was clarified by centrifugation (Sorvall RC-5B Superspeed Centrifuge) at 15,000 rpm (68,040x *g*) at 4 °C for 45 min. The supernatant was incubated with Ni-NTA agarose beads (Qiagen) for 1 h. The Ni-NTA beads were washed with 20 column volumes (CV) of lysis buffer, 10 CV of wash buffer (50 mM HEPES-NaOH, pH 7.5, 1M NaCl, 5 mM MgCl_2_, 20 mM imidazole, 1mM CHAPS, 5mM 2-mercaptoethanol, 10% (v/v) glycerol), and 10 CV of lysis buffer again before the protein was eluted with 5 CV elution buffer (50 mM HEPES-NaOH, pH 7.5, 500 mM NaCl, 5 mM MgCl_2_, 500 mM imidazole, 1mM CHAPS, 5mM 2-mercaptoethanol, and 10% (v/v) glycerol). The sample was then dialyzed into lysis buffer containing Ulp1 protease to remove the N-terminal 6x-His-SUMO tag overnight at 4 °C. The dialyzed protein was concentrated using a 30 kDa MWCO concentrator (Millipore) and was further purified by size-exclusion chromatography on a Superose 6 Increase 10/300 GL column (Cytiva) equilibrated with purification buffer (20 mM HEPES-NaOH, pH 7.5, 140 mM NaCl, 10 mM KCl, 1mM MgCl_2_, and 1 mM DTT). A typical size-exclusion profile for WT LACTB consisted of a large peak containing LACTB filaments eluting within the void volume (V_0_) of the column. Fractions of LACTB were analyzed by SDS-PAGE. Freshly purified samples were used for cryoEM grid preparation and *in vitro* substrate binding assays.

### Negative-stain electron microscopy

The LACTB assembly was purified as described above. Size-exclusion peak fractions corresponding to LACTB were diluted to ∼2.0 μM for negative-stain transmission electron microscope (TEM). Samples were prepared by applying 4 μl of purified LACTB applied onto glow-discharged carbon-coated copper grids (400 mesh) and negatively stained with 0.75% (w/v) uranyl formate (UF) following an established protocol [67]. Briefly, the samples were incubated on the grid for 1 min and then blotted off with filter paper. Subsequently, samples on grids were stained with UF for 45 s and air dried for 2 min at room temperature. The grids were imaged using a Tecnai T12 Spirit LaB6 filament TEM (FEI) equipped with an AMT 2k x 2k side-mounted CCD camera and operated at a voltage of 100 kV. The micrographs were collected at a nominal magnification of 120,000x at the specimen level with a calibrated pixel size of 5.28 Å per pixel.

### CryoEM sample preparation and data acquisition

Prior to cryoEM grid preparation, an aliquot of purified LACTB was buffer exchanged to a freezing buffer (20 mM HEPES-NaOH, pH 7.5, 140 mM NaCl, 10 mM KCl, 1 mM MgCl_2_, 1mM DTT, 1% glycerol, and 1 mM CHAPS) by centrifugation at 1000x *g* for 2 minutes with a micro-spin desalting column (Thermo Scientific Zeba) at 4 °C. Aliquots of 4 μl of protein samples (0.3 mg ml^-1^ or ∼6 μM) were applied to glow-discharged R1.2/1.3 200 Cu mesh grids (Quantifoil). After 30 s, the grids were blotted for 3 seconds and plunged into liquid ethane using a Leica EM GP2 Automatic Plunge Freezer operated at 10 °C and 90% humidity. Micrographs were acquired using a Titan Krios TEM (FEI) operated at 300 kV and equipped with a K3 Summit direct electron detector (Gatan). Images of frozen-hydrated filaments of LACTB were collected in counting mode at a nominal magnification of 29,000x corresponding to an image pixel size of 0.8211 Å on the specimen level, applying a defocus range of -0.5 to -1.5 μm. A total of 7643 dose-fractionated movies were recorded with the SerialEM software [68] multishot method over nine neighboring holes (3 x 3). The dose rate was 1.05 electrons/Å^2^/frame, with a frame rate of 0.07 s, resulting in a total exposure time of 3.985 s and a total accumulated dose of 59.58 electrons per Å^2^ (57 frames). CryoEM data collection parameters are summarized in Table 1.

### Image analysis and 3D reconstruction

The alignment and dose-weighted summation of raw movie stacks were carried out using MotionCor2 [69]. Contrast transfer function (CTF) estimations were performed on aligned, non-dose-weighted micrographs using CTFFIND4 [70]. All subsequent processing steps were performed using Relion 3.1 [71,72]. A total of 34,636 filament coordinates were manually picked and filament segments with 90% overlap were extracted by using a box size of 320 pixels. All segments (430,592) were then subjected to multiple rounds of reference-free two-dimensional (2D) classification with 100 classes to identify structurally homogeneous subsets. After 2D classification, a featureless cylinder-shaped electron density map with a 160 nm diameter was generated in Relion and used as an initial reference for 3D auto-refinement with C1 symmetry. The resulting refined map showed clear secondary structure features and was used to estimate the helical parameters. Using this 4.3 Å reconstruction, an initial 3D classification was performed with C1 symmetry, and the segments from the best 3D classes (248,548 segments) were selected for high-resolution 3D auto-refinement. Subsequently, these segments were 3D auto-refined with a soft mask, central Z length of 45% of the box size (helical_z_percentage=0.45), and imposing starting helical parameters of 21.59 Å rise and 48.91° twist, yielding a 3D reconstruction with an overall resolution of 3.7 Å. To improve this density map, the final particle segments were subjected to iterative rounds of focused 3D auto-refinement, CTF refinement, and Bayesian polishing to obtain a final 3D reconstruction of a right-handed helical assembly at 3.1 Å resolution. The helical parameters converged to a rise of 21.63 Å and a twist of 48.25° per subunit. A reconstruction without imposing symmetry was also generated at 3.1 Å resolution after focus refinement using a similar mask on the filament. The final reported resolutions are based on the gold-standard Fourier shell correlation (FSC) = 0.143 criterion. Maps were sharpened by applying a negative *B* factor as determined by the Relion post-processing program. The local resolution estimation was calculated by ResMap using half-map reconstructions [73].

### Model building and refinement

Model building was carried out using the 3.1 Å cryoEM map of the human LACTB filament. The Alphafold [74] predicted structure of monomeric LACTB was used as an initial model for atomic model building. The model was manually fitted into the cryoEM density map using UCSF Chimera [75] and then rigid body refined using the real-space refinement procedure implemented in the program PHENIX [76]. Atomic coordinates of the human LACTB (residues 97-547) filament were manually rebuilt using the program Coot [77] and the final models were iteratively refined against the cryoEM density map. The real space-refinement of models with global minimization, local grid search, secondary structure, and geometric restraints applied was performed using the software PHENIX [76]. Secondary structure elements and side chains for the majority of the residues were unambiguously resolved. Each residue was manually inspected, and correction of the refined coordinates was performed by applying torsion, planar peptide, and Ramachandran restraints during manual rebuilding in Coot [77].

### Validation and structural analysis

The stereochemistry and geometry of the final model was evaluated by MolProbity [78]. CryoEM maps and atomic models were analyzed, and figures were generated using UCSF ChimeraX [79], UCSF Chimera [75], and PyMOL (The PyMOL Molecular Graphics System, Version 2.0 Schrödinger, LLC). Negative-stain TEM images were analyzed using FIJI [80]. Surface potential renderings were generated using the APBS plugin in PyMOL [81]. Structural and chemical properties of macromolecular interfaces and surfaces were analyzed by the PISA server [82]. The fold conservation analysis was performed with the Dali server [83] and structural comparisons to the PBP-βL family of enzymes and RMSD calculations were carried out using the CLICK server [84]. Sequence alignment of the LACTB homologs as well as the PBP-βL family of enzymes was generated using Clustal Omega [85] and sequence conservation figures were prepared by using the Clustal Omega alignment file and LACTB model in ESPript server 3.0 [86]. Schematic diagrams were generated by using BioRender.com. The statistics of the cryoEM helical reconstruction and model refinement were summarized in Table 1.

### LACTB *in vitro* substrate assay

To determine the catalytic activity of the WT and mutant LACTB, the cleavage of the fluorogenic Ac-YVAD-AMC peptide (Enzo Life Sciences) was measured at an excitation wavelength of 380 nm and an emission wavelength of 460 nm using a CLARIOstar microplate reader. All assays were performed at room temperature and the total reaction volumes were 20 μl. Briefly, 10 μl of purified LACTB protein was mixed with a reaction buffer containing 20 mM HEPES-NaOH, pH 7.5, 140 mM NaCl, 10 mM KCl, 1 mM MgCl_2_, 1 mM DTT, and 0.01% NP-40 detergent (AmericanBio) to generate a final protein concentration of 12-15 nM. Reactions were initiated by adding 100 μM concentrations of the substrate Ac-YVAD-AMC (dissolved in DMSO and prediluted to 200 μM in reaction buffer) to the enzyme in a 384 well black polystyrene assay plate (Costar 3916). The reactions were run in at least three independent replicates for 1,200 s and fluorescence was measured every 10 s. The raw data were averaged to determine reaction rates from the linear part of the curve and include standard deviation. To determine the catalytic activity of the WT LACTB in the presence of lipid vesicles, freshly purified protein (1.5-2 μM) was incubated with 0.5 mg ml^-1^ unilamellar liposomes in three independent experiments for 4 h at room temperature. Then, the catalytic activity of LACTB was measured as described above to quantify the reaction rate of WT LACTB in the presence of liposomes.

### H_2_O_2_ treatment of LACTB filaments

An aliquot of purified LACTB was diluted to 0.15 mg ml^-1^. Aliquots of 10 μl of the protein sample were incubated with 5 μl of varying concentrations of H_2_O_2_ for 30 minutes. The final concentrations were 0.01% H_2_O_2_ (v/v), 0.025%, 0.05%, 0.1%, 0.25%, 0.5%, 0.75%, and 1.0%. The final concentration of protein was 0.1 mg ml^-1^. The samples were analyzed by SDS PAGE using a 4-15% precast gel (BioRad) at 100 V. The 0.1%, 0.5%, and 1.0% H_2_O_2_ (v/v) samples were stained on glow-discharged carbon-coated copper grids with UF as described above and visualized with a Tecnai T12 Spirit TEM (FEI) operated at a voltage of 100 kV. The micrographs were collected at a nominal magnification of 120,000x at the specimen level with a calibrated pixel size of 5.28 Å per pixel.

### Preparation of lipid vesicles and LACTB membrane binding reactions

To determine the membrane binding activity of human LACTB, the lipid composition of the mitochondrial inner membrane was used to prepare vesicles as described previously [87]. The lipid stock solutions were resuspended in chloroform, methanol, and water mixture (20:9:1, (v/v/v)) and stored at -20°C. 1-palmitoyl-2-oleoyl-glycero-3-phosphocholine (POPC), 1-palmitoyl-2-oleoyl-sn-glycero-3-phosphoethanolamine (POPE), L-α-lysophosphatidylinositol (Soy Lyso PI), 1’,3’-bis[1,2-dioleoyl-sn-glycero-3-phospho]-glycerol (cardiolipin (18:1)_4_), D-galactosyl-ß-1,1’ N-nervonoyl-D-erythro-sphingosine (C24:1 Galactosyl(ß) Ceramide, GalCer) (Avanti Polar Lipids) were mixed in a molar ratio of POPC:POPE:PI:CL; 45:22:8:25, POPC:POPE:PI; 70:22:8, POPC:POPE; 78:22, POPC; 100, and GalCer:CL; 80:20. The lipid mixture was then dried as a thin film in a glass tube under a gentle stream of nitrogen and incubated in a vacuum desiccator for 4 h to remove excess solvent. Following dehydration, the lipid film was resuspended for 30 min at room temperature in a liposome buffer containing 20mM HEPES-NaOH, pH 7.5, and 150mM NaCl. The suspension was extruded through polycarbonate membranes with a 50 nm pore diameter using Avanti Mini Extruder (Avanti Polar Lipids) to obtain unilamellar liposomes. 20 μl aliquots of rehydrated lipids were flash-frozen in liquid nitrogen and stored at -80°C. The nanotube mixtures were rehydrated in liposome buffer and incubated at room temperature for 15 min with intermittent mixing. The suspended mixture of lipid nanotubes was briefly vortexed and sonicated for three rounds (5 min per round) in a bath sonicator at 50°C. The lipids were immediately used in the binding assays after cooling at room temperature.

For liposome binding assays, a range of 1.5-2 μM of freshly purified WT LACTB was incubated with 0.5 mg ml^-1^ unilamellar liposomes or the liposome reaction buffer (20 mM HEPES-NaOH, pH 7.5, 140 mM NaCl, 10 mM KCl, 1 mM MgCl_2_, 1 mM DTT) for 4 h at room temperature. Subsequently, the reaction mixture was applied onto glow-discharged carbon-coated copper grids and negatively stained with UF. The membrane binding activity of LACTB filaments was assessed by negative-stain TEM. Images were collected using a Tecnai T12 Spirit TEM (FEI) equipped with an AMT 2k x 2k side-mounted CCD camera as described above. Negative stain images were recorded at a nominal magnification of 120,000x at the specimen level with a calibrated pixel size of 5.28 Å per pixel. A total of 1000 filaments were counted from randomized blinded images to eliminate bias. The total percentage of filaments interacting with one or more liposomes (binding and tethering, respectively) was determined and the s.e.m. for each sample was calculated.

### Statistical analysis

Statistical significance was determined using GraphPad Prism 9 software. Liposome tethering experiments and related enzymatic activity measurements were analyzed by unpaired two-tailed Welch’s t-test and unpaired two-tailed Student’s t-test, respectively. The values P < 0.05 were considered statistically significant.

### Data Accessibility

All of the 3D cryoEM data that support the findings of this study have been deposited in Electron Microscopy Data Bank with the accession code: EMDB-26595. The model coordinates have been deposited in the Protein Data Bank under accession code PDB ID: 7ULW.

## Acknowledgments

We thank the current members of the Aydin laboratory for helpful discussions and assistance on this project. We especially thank Charles Moe at the BioChemistry Krios Electron Microscopy (BioKEM) Facility of the University of Colorado Boulder for his support with data collection and EM infrastructure; Garry Morgan and Courtney Ozzello at the EM Services Core Facility of the University of Colorado Boulder for electron microscopy training and support; the Shared Instrument Pool (SIP) core facility (RRID: SCR_018986) of the Department of Biochemistry at the University of Colorado Boulder for the use of the shared research instrumentation infrastructure; Dr. Annette Erbse for assistance with biophysical instruments and support; Dr. Chris Ebmeier and the Mass Spectrometry Core Facility of the University of Colorado Boulder, which is supported by NIH Grants S10OD025267 and S10RR026641 grants, for mass spectrometry analysis; Dr. Kelly Zuccaro for assistance with computer infrastructure and liposome preparation. We also thank Professor Karolin Luger for her support and for kindly sharing the equipment for the fluorescence-based activity assays. The results shown here are in part based upon data generated by the TCGA Research Network: https://www.cancer.gov/tcga.

## Competing interests

The authors declare no competing interests.

**Correspondence and requests for materials** should be addressed to H.A.

## Supporting Information

**S1 Fig.**
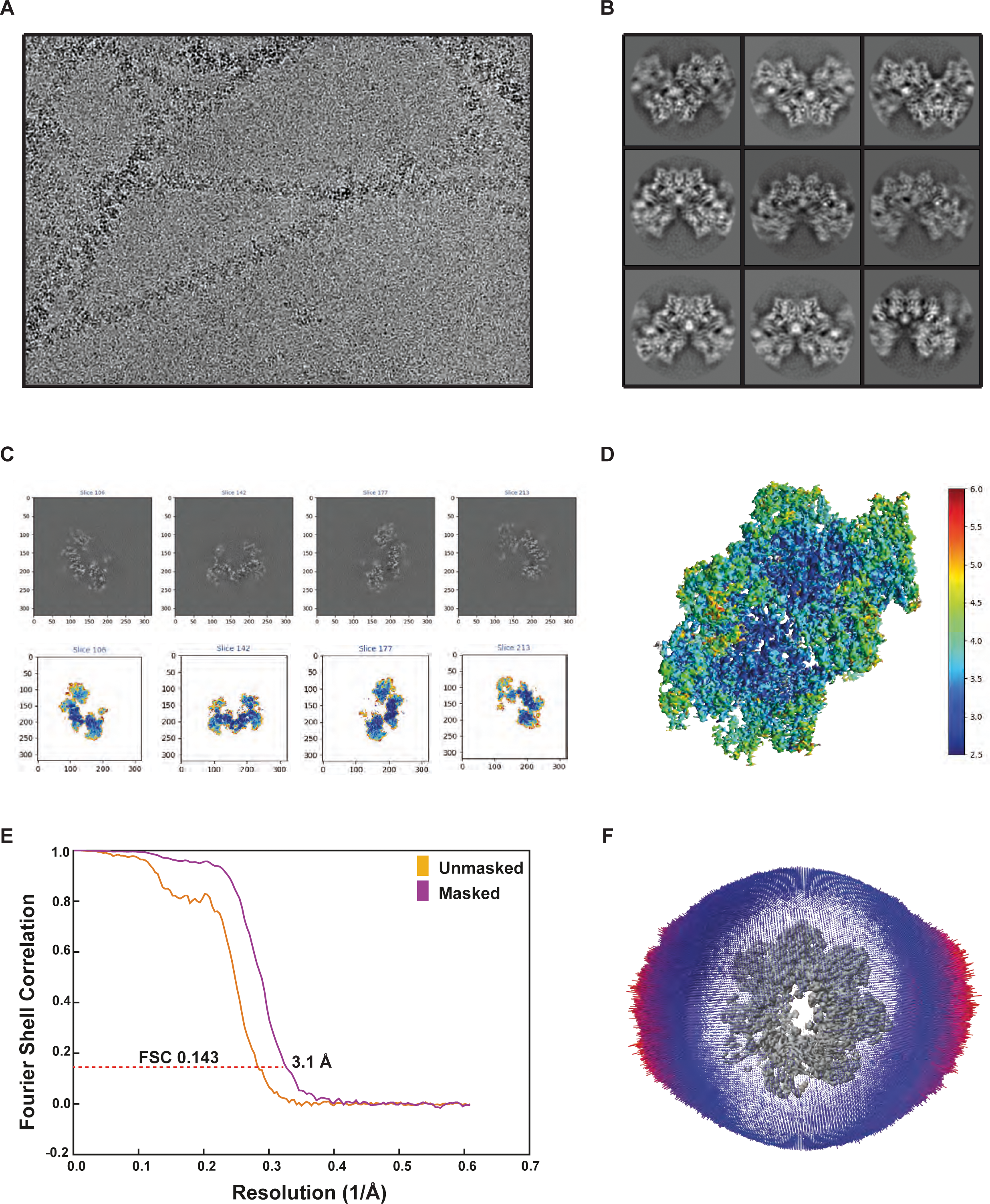
CryoEM raw data and validation of the human LACTB filament. (**A**) A representative cryoEM micrograph and (**B**) 2D class averages of the human LACTB filament. (**C**) Slices through the unsharpened density map of the LACTB filament are shown in the top view. (**D**) CryoEM map of the human LACTB filament is colored according to the local resolution estimation calculated by ResMap. (**E**) Fourier Shell Correlation (FSC) curve shows 3.1 Å global resolution (masked in purple and unmasked in orange) with the gold standard criteria (FSC=0.143). (**F**) Euler angle distribution of the segments used in the final 3D reconstruction.

**S2 Fig.**
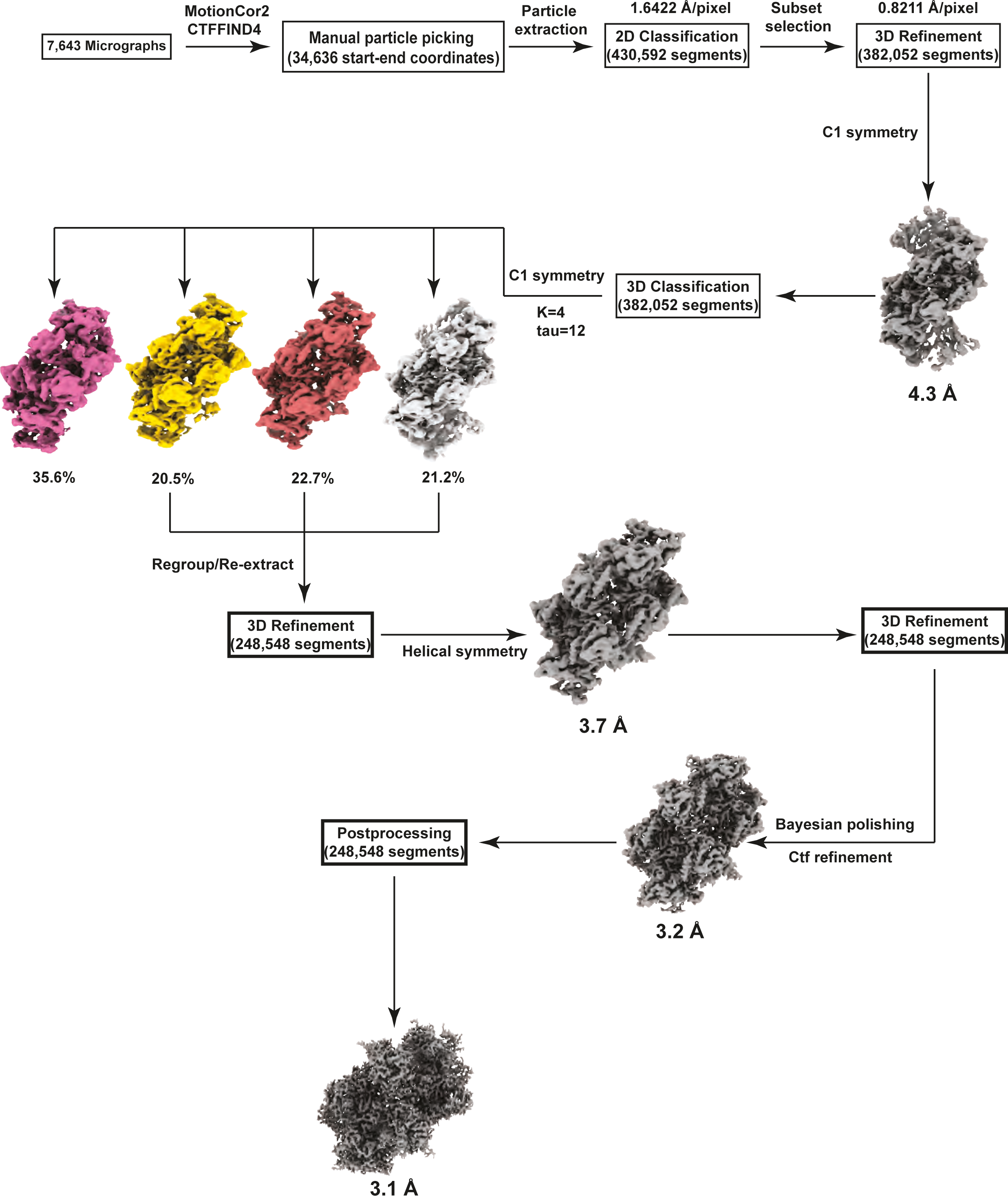
LACTB filament data processing strategy. Flow chart summarizing cryoEM image-processing workflow, including 3D classification and 3D auto-refinement, CTF refinement, and Bayesian polishing.

**S3 Fig.**
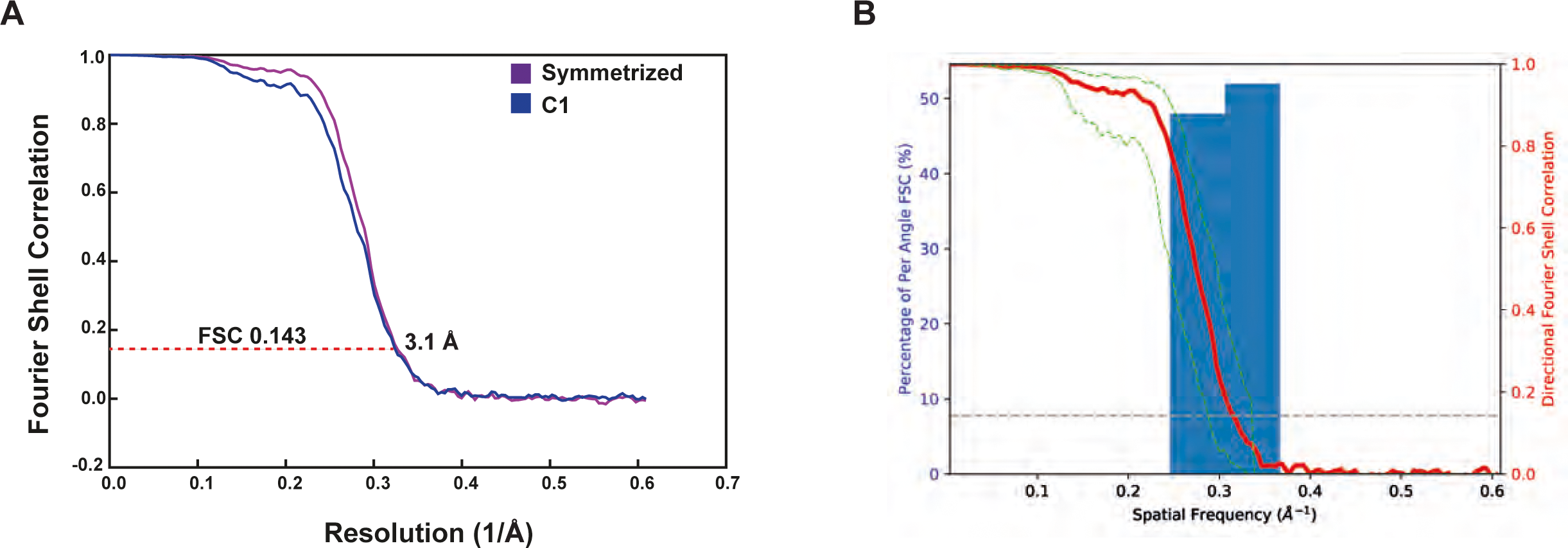
CryoEM map quality of the LACTB filament. (**A**) Fourier Shell Correlation (FSC) curves for the final helical symmetry imposed (purple line) and C1 symmetry (blue line) 3D reconstructions. (**B**) 3D Fourier Shell Correlation of the final cryoEM map.

**S4 Fig.**
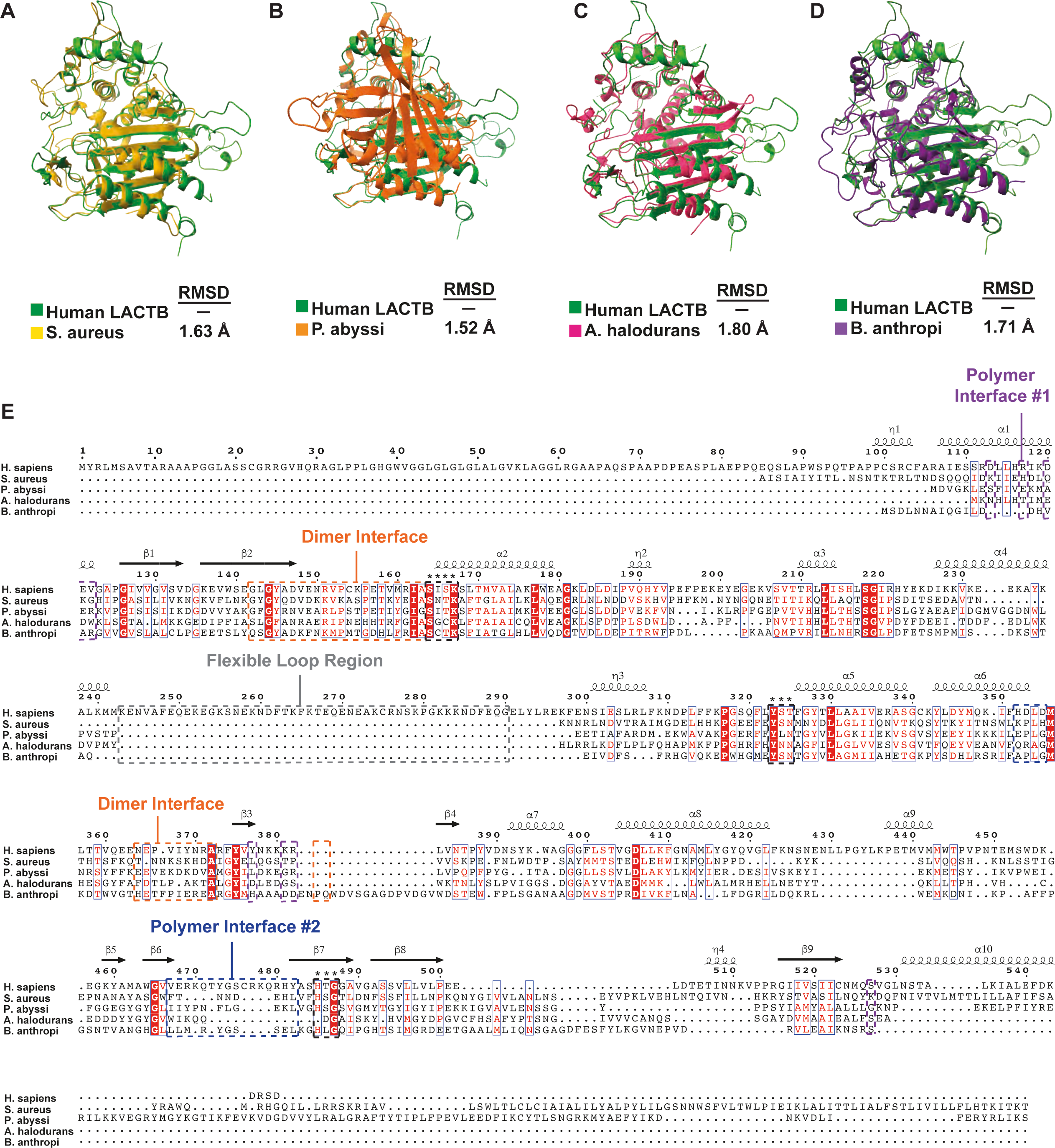
Comparison of the LACTB monomer with different structures of homologous proteins and sequence alignment with bacterial PBP-βL family of enzymes. (**A-D**), Comparison of human LACTB monomer (green) with *S. aureus* ClbP protein (yellow, PDB ID: 4GDN), *P. abyssi* Pab87 peptidase (orange, PDB ID: 2QMI), *A. halodurans* penicillin-binding protein (pink, PDB ID: 3TG9), and *B. anthropi* D-amino-acid amidase (purple, PDB ID: 2EFU). The root-mean-square deviation of each structure relative to the LACTB monomer is shown at the bottom. While most β-strands align well between human and bacterial proteins, the surrounding helices and interface-forming loops show significant differences between these structures. (**E**) Sequence alignment of LACTB homologs and orthologs. Residues that form the dimerization interface and polymerization interfaces 1 and 2 are highlighted with orange, purple and blue boxes, respectively. Catalytic site residues are highlighted with an asterisk. Secondary structure elements shown above the alignment are generated from the human LACTB structure. Sequences were aligned using Homo Sapiens LACTB (Uniprot ID: P83111), *S. aureus* ClbP (Uniprot ID: Q7A3Q5), *P. abyssi* Pab87 peptidase (Uniprot ID: Q9V2D6), *A. halodurans* penicillin-binding protein (Uniprot ID: Q9KAM0), and *B. anthropi* D-amino-acid amidase (Uniprot ID: Q9LCC8).

**S5 Fig.**
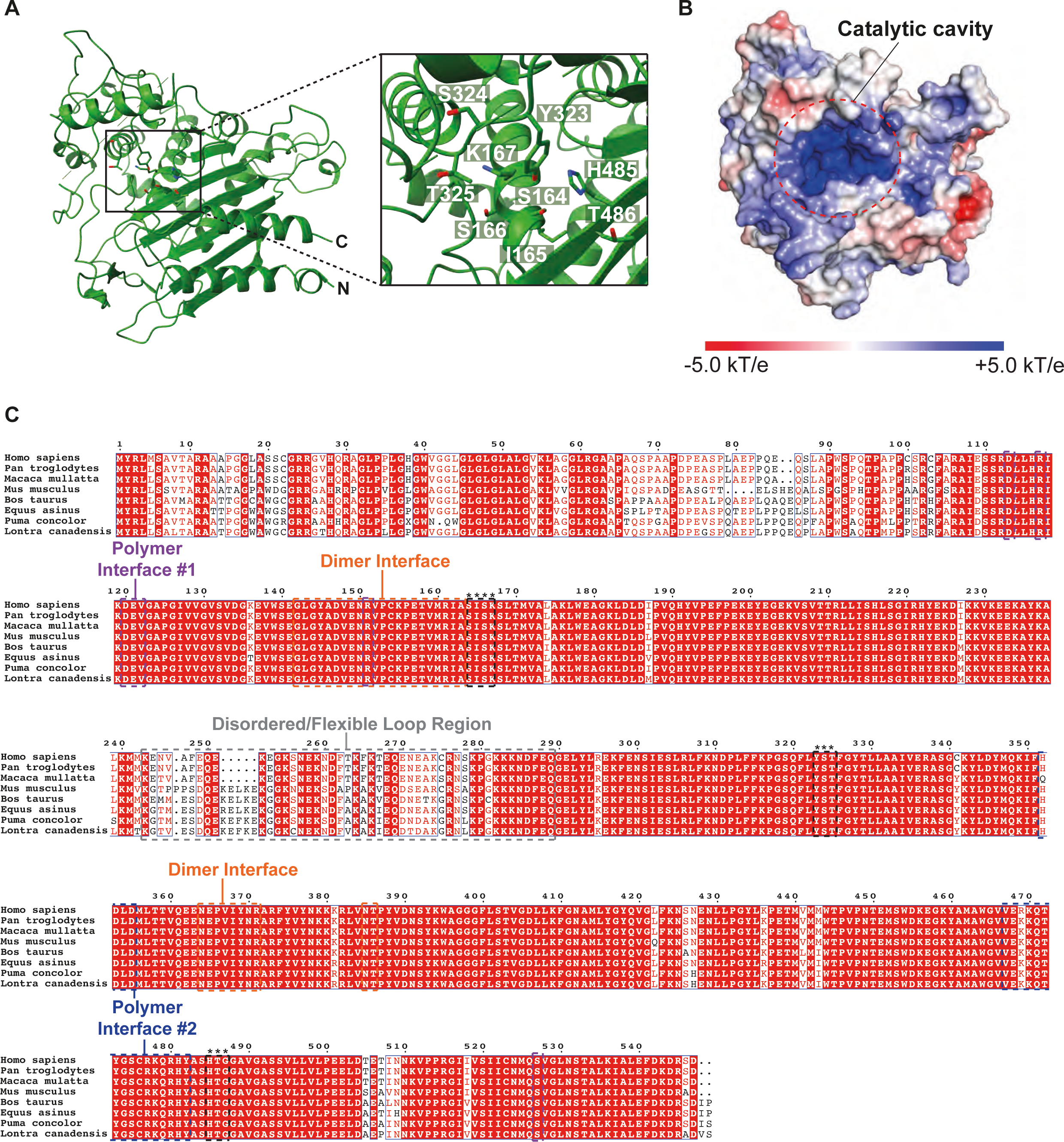
Structure of LACTB catalytic site and multiple sequence alignment of mammalian LACTB homologs. (**A**) Active site and ligand recognition of human LACTB. Ribbon diagram of a monomer is displayed in green and residues forming the catalytic site are shown as sticks. Boxed zoom images highlight the positions of the conserved residues that contribute to catalytic site formation. (**B**) Surface electrostatic potential representation of the LACTB monomer. Positive and negative electrostatic potentials are shown in blue and red, respectively. The substrate binding cavity is highlighted with a red circle. (**C**) Sequence alignment of LACTB homologs. Residues that form the dimerization interface and polymerization interfaces 1 and 2 are highlighted with orange, purple and blue boxes, respectively. Catalytic site residues are highlighted with an asterisk. Secondary structure elements shown above the alignment are generated from the human LACTB structure. LACTB sequences from Homo sapiens (human; Uniprot ID: P83111), Pan troglodytes (chimpanzee; Uniprot ID: K7CYM3), Macaca mulatta (rhesus macaque; Uniprot ID: F7EXQ6), Mus musculus (mouse; Uniprot ID: Q9EP89), Bos taurus (cow; Uniprot ID: P83095), Equus asinus (donkey; Uniprot ID: UPI001D03C0DD), Puma concolor (mountain lion; Uniprot ID: A0A6P6HFT1), and Lontra canadensis (river otter; Uniprot ID: UPI0013F34634) are aligned.

**S6 Fig.**
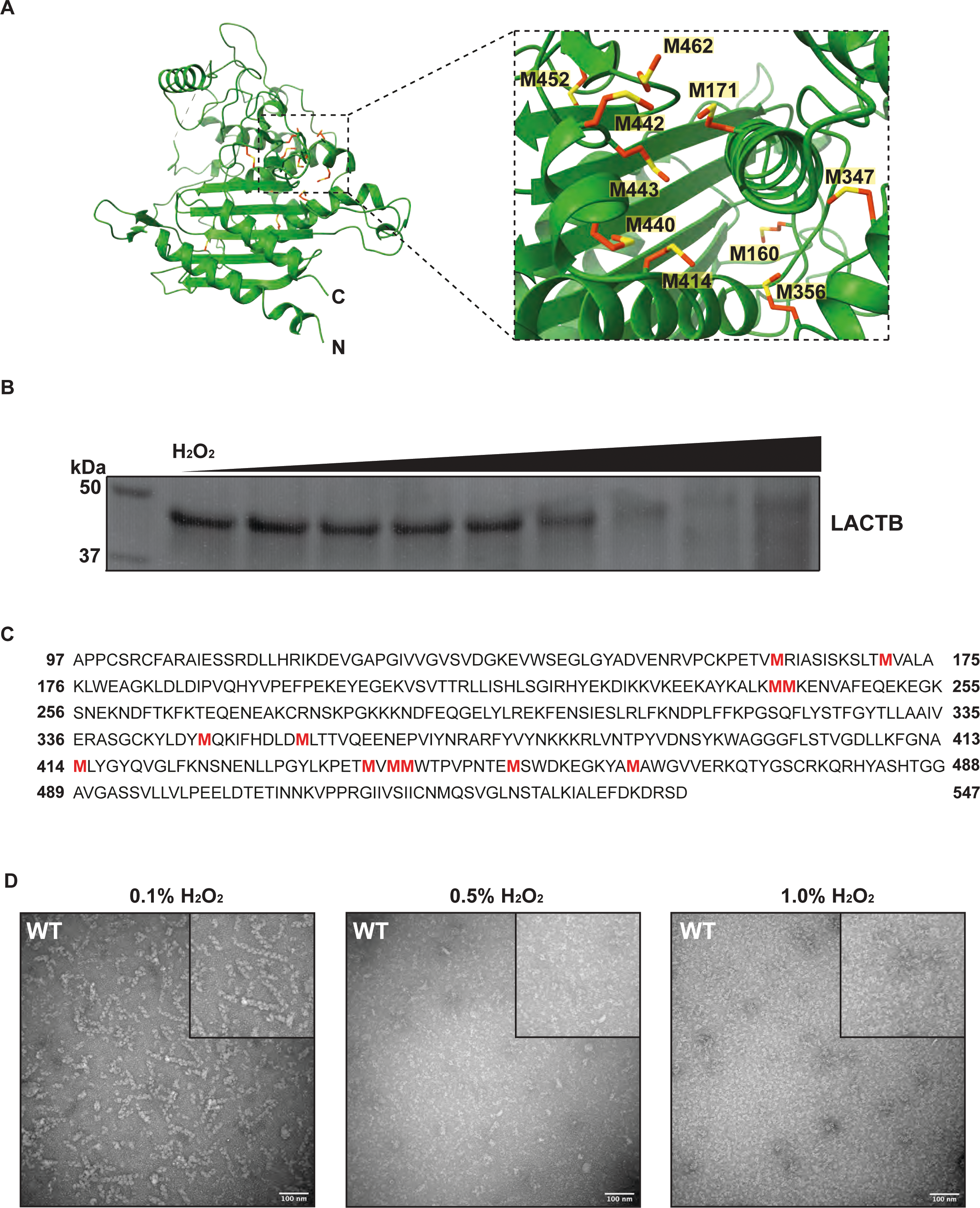
SDS-PAGE and negative-stain TEM analyses of LACTB filaments in the presence of H_2_O_2_. (**A**) Ribbon diagram of the LACTB monomer is displayed in green, and residues forming a methionine cluster are shown as sticks. Boxed zoomed in image highlights the positions of methionine residues which are colored orange and yellow. (**B**) Coomassie blue-stained SDS-PAGE gel of WT LACTB protein treated with increasing concentrations (0.1%, 0.5%, and 1.0%) of H_2_O_2_. Indicated concentrations of H_2_O_2_ were incubated with purified LACTB for 30 min before SDS-PAGE analysis. The left lane is the molecular weight standards, and their positions are indicated. (**C**) Sequence of WT LACTB construct (residues 97-547) with confirmation of methionine residues oxidized upon exposure to H_2_O_2_ (shown in red) by liquid chromatography with tandem mass spectrometry (LC-MS/MS). The underlying data for (**C**) is provided in S2 Data. (**D**) Representative negative-stain TEM micrographs of WT LACTB treated with varying concentrations of H_2_O_2_ for 30 min before grid preparation. Scale bars, 100 nm.

**S7 Fig.**
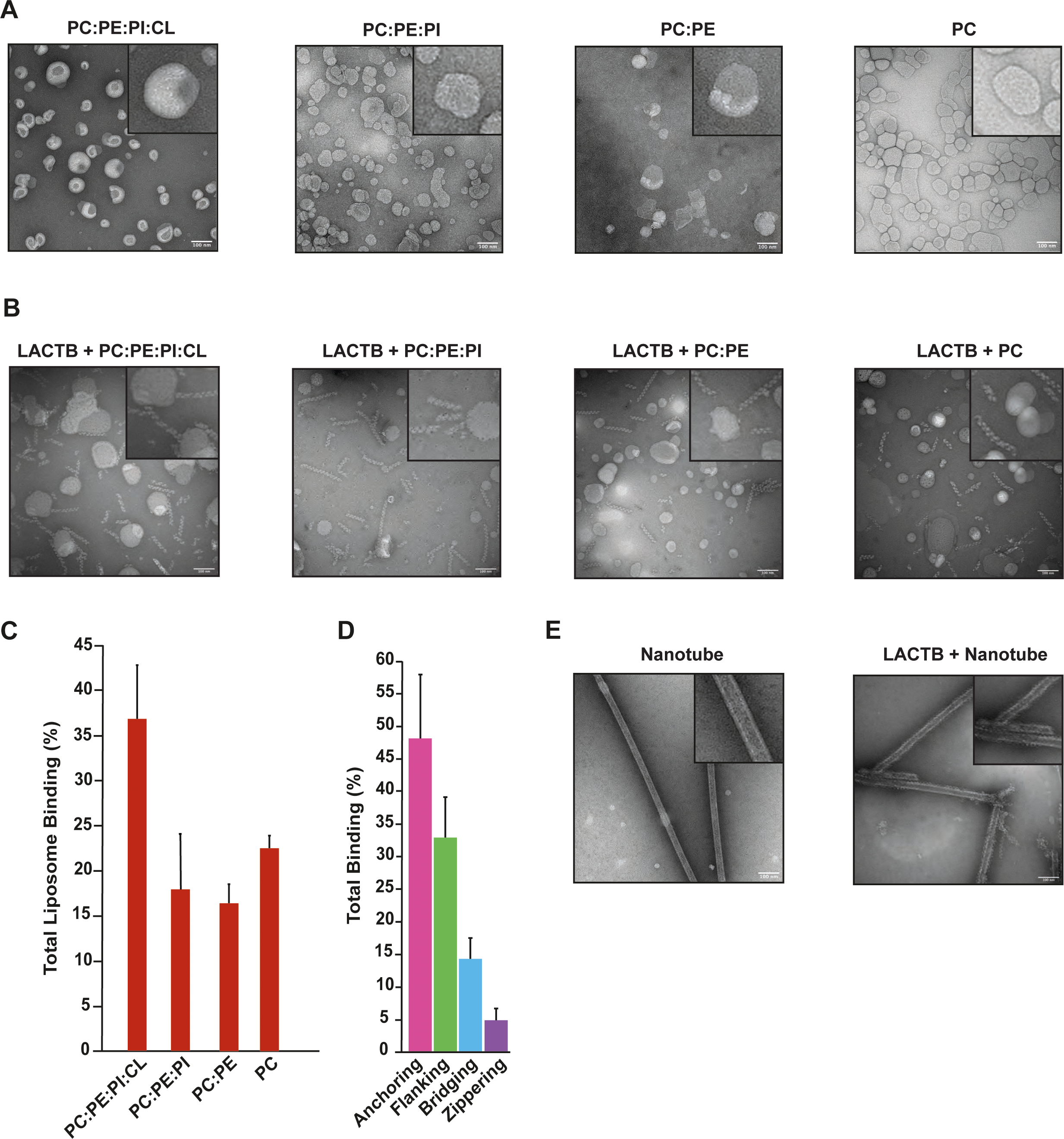
Structural characterization of LACTB filaments with liposomes of different compositions and nanotubes. (**A**) Negative stain images of liposomes with different compositions. From left to right: PC:PE:PI:CL, PC:PE:PI, PC:PE, PC. (**B**) Negative stain images of WT LACTB in the presence of liposomes with different compositions in the same order as (**A**). (**C**) Total liposome binding activity of WT LACTB with liposomes of different compositions. Total liposome binding was quantified by counting 1000 filaments from at least three independent reconstitution assays and calculating the total percentage of filament binding. Bars represent the mean of at least three independent reconstitution assays and error bars indicate the s.e.m. (**D**) Percentage of LACTB filaments bound to liposomes by interaction type. Bars represent the mean of at least three independent reconstitution assays and error bars indicate the s.e.m. (**E**) Negative stain images of empty nanotubes (left) and WT LACTB filaments in the presence of nanotubes (right). The source data for (**C**) and (**D**) are provided in sheets S7C and S7D Fig in S1 Data.

**S8 Fig.**
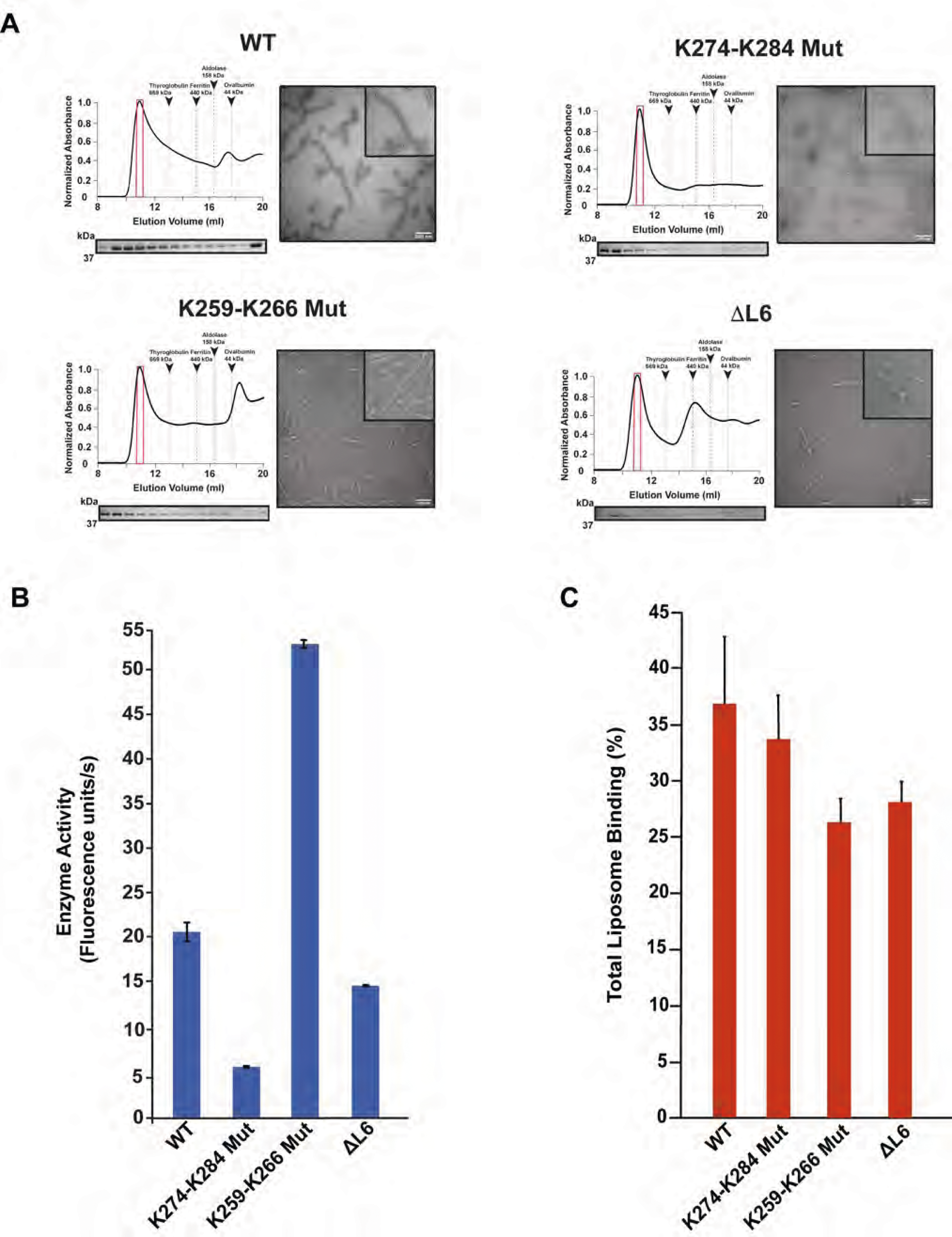
Structural characterization of the lipid binding region of LACTB. (**A**) Size exclusion chromatography (SEC) elution profiles of WT and mutant LACTB. SEC elution profiles begin with the void volume (V_0_) of the column. Fractions were analyzed by SDS-PAGE and peak fractions used in negative-stain analyses are highlighted in red boxes. Negative-stain TEM images demonstrate the polymerization activity of WT and mutant proteins. Scale bars, 100 nm. (**B**) Catalytic activity of key residues involved in lipid binding. Enzyme activity was determined *in vitro* by using a fluorescently labeled substrate and three independent experiments were performed for each sample. Bars represent the mean of at least three independent experiments and error bars represent the standard deviation. (**C**) Total liposome binding activity of WT LACTB and key residues involved in lipid binding. Total liposome binding was quantified by counting 1000 filaments from at least three independent reconstitution assays and calculating the total percentage of filament binding. Bars represent the mean of at least three independent reconstitution assays and error bars indicate the s.e.m. The underlying data in (**B**) and (**C**) are provided in sheets S8B and S8C Fig in S1 data.

**S9 Fig.**
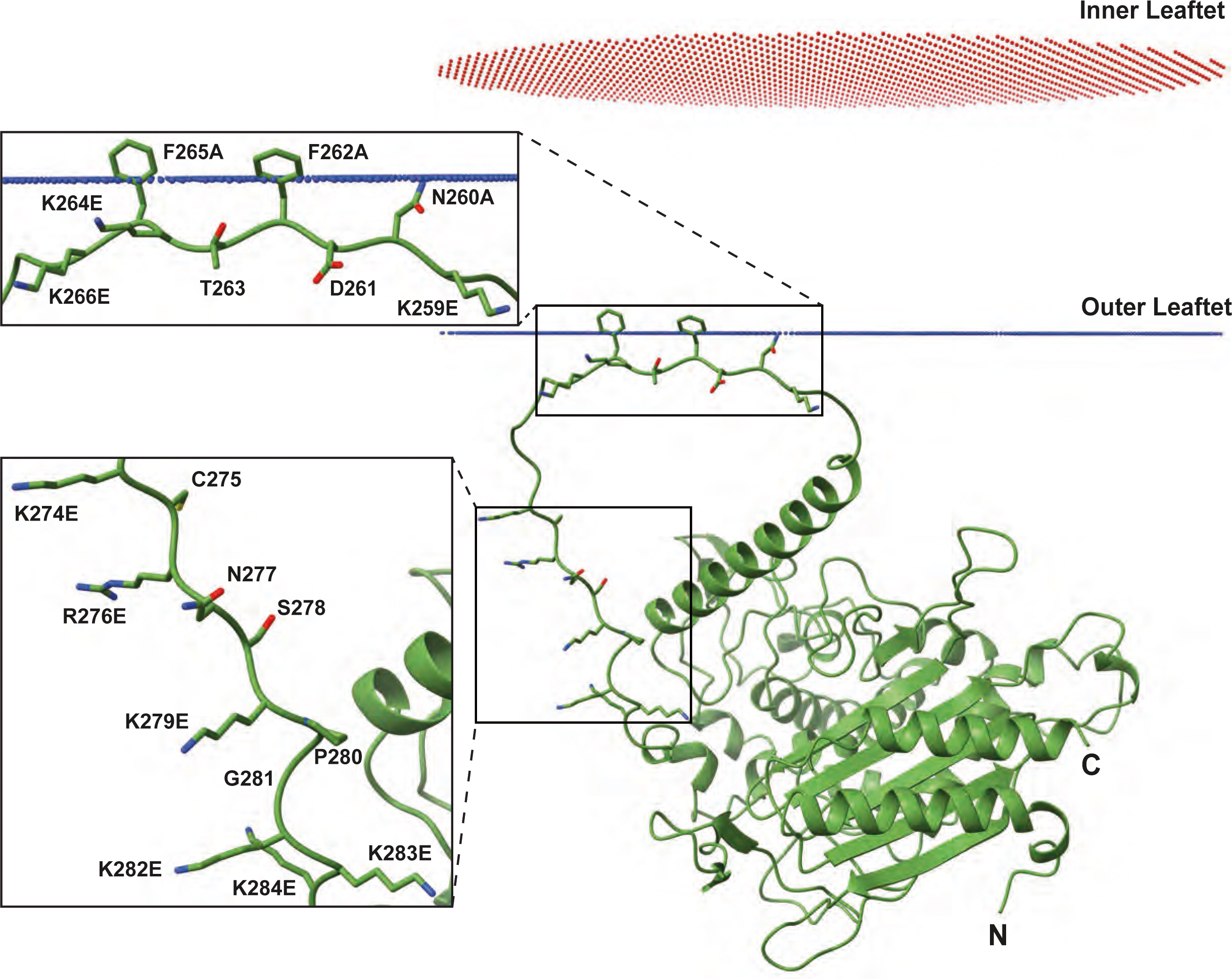
Computational modeling of LACTB lipid binding motifs to membranes. The monomeric structure of WT human LACTB was obtained from AlphaFold and the rotational and translation position of the protein was calculated in the presence of membranes, mimicking the lipid composition of the mitochondrial IM using the PPM Web Server. Boxed zoomed in images highlight the predicted structures of ^259^KNxFxKFK^266^ and ^274^KxRxxKxxKKK^284^ motifs within the flexible loop region (L6) of human LACTB.

**S10 Fig.**
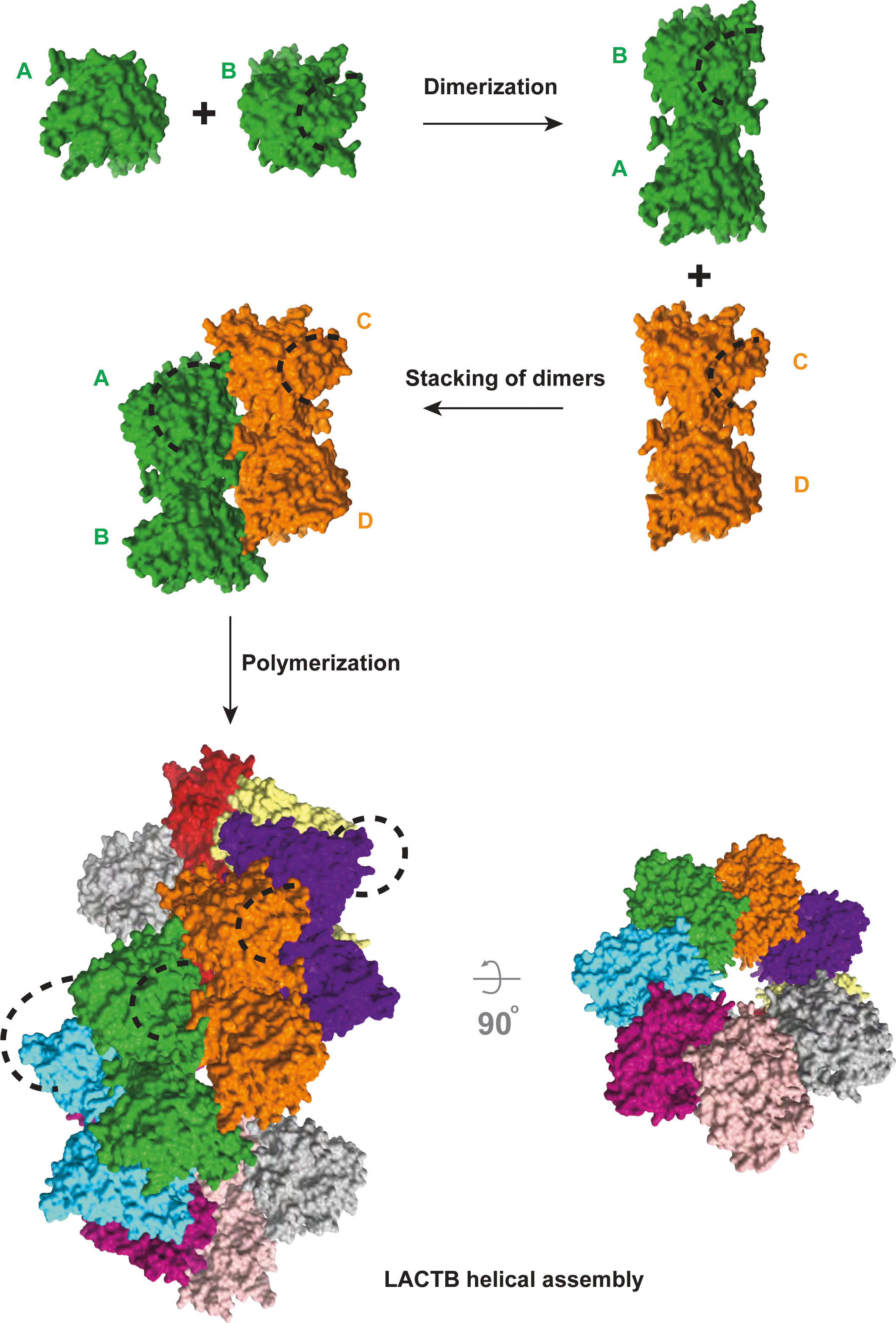
Mechanistic model describing LACTB assembly mechanism within mitochondria. After being imported to mitochondrial IMS, LACTB protomers assemble into antiparallel dimers. Subsequently, LACTB dimers interact through two polymerization interfaces, which allow the stacking of antiparallel dimers in a helical fashion and lead to the polymerization into micron-scale helical assemblies. Upon filament formation, LACTB gains catalytic activity and filament elongation increases the catalytic efficiency of the enzyme.

**S1 Table.** Disease-associated missense mutations in human LACTB. Mutations associated with various cancers were compiled from the Catalogue of Somatic Mutations in Cancer (COSMIC), the Genome Aggregation Database (gnomAD), and The Cancer Genome Atlas Program (TCGA).

| **Protein change** | **Clinical classification** | **Cancer type** |
| --- | --- | --- |
| H116Q | Neutral | Thyroid carcinoma |
| E121K | Pathogenic | Breast |
| V148F | Pathogenic | Kidney |
| E149Q | Pathogenic | Esophagus |
| R151S, R151H | Likely pathogenic | Uterus |
| E363K | Pathogenic | Pancreatic |
| R371K | Pathogenic | Lung |
| A372T | Pathogenic | Uterus |
| K380N | Pathogenic | Cervical |
| R382L, R382C | Pathogenic | Oral, uterus |
| E457K | Pathogenic | Bladder |
| E468G | Pathogenic | Glioma |
| R469K | Pathogenic | Colon, breast |
| T472K, T472M | Neutral | Melanoma, colon |
| R480W, R480Q, R480L | Pathogenic | Lung, colon, liver |
| Y482H | Pathogenic | Melanoma |

Mutations associated with cancer were compiled from the Catalogue of Somatic Mutations In Cancer (COSMIC), the Genome Aggregation Database (gnomAD), and The Cancer Genome Atlas Program (TCGA).

**S1 Data. The numerical values underlying Figs 1F and 3D and 4E and 5G-I and S7C-D and S8B-C**. (XLSX). Sheets show the values underlying the data displayed in all figures and includes an explanation of all data. Sheet Fig 1F shows the raw data from *in vitro* substrate fluorescence assays for WT LACTB underlying Fig 1F. Sheet Fig 3D details the raw data from *in vitro* substrate fluorescence assays for LACTB mutants underlying Fig 3D. Sheet Fig 4E describes the raw data from *in vitro* substrate fluorescence assays for LACTB mutants underlying Fig 4E. Sheet Fig 5G shows the raw data from *in vitro* substrate fluorescence assays in the presence of liposomes underlying Fig 5G. Sheet Fig 5H describes the raw data from LACTB membrane binding assays based on membrane composition underlying Fig 5H. Sheet Fig 5I reports the raw data from LACTB membrane binding assays for LACTB mutants underlying Fig 5I. Sheet S7C Fig represents the raw data from LACTB membrane binding assays based on membrane composition underlying S7C Fig. Sheet S7D Fig shows the raw data from LACTB membrane binding assays for WT LACTB based on binding classification underlying S7D Fig. Sheet S8B Fig details the raw data from *in vitro* substrate fluorescence assays for LACTB mutants underlying S8B Fig. Sheet S8C Fig describes the raw data from LACTB membrane binding assays for LACTB mutants underlying S8C Fig.

**S2 Data. Source data of liquid chromatography with tandem mass spectrometry (LC-MS/MS) in S6C Fig.** (XLSX). Parameters sheet shows the experimental parameters for LC-MS/MS experiments of WT LACTB in the presence of H_2_O_2_. Summary sheet describes the summary of experimental data for LC-MS/MS experiments. Evidence sheet shows protein sequence information for LC-MS/MS experiments. Protein Groups sheet details protein identity information for LC-MS/MS experiments.

**S1 Raw Images. Uncropped images of SDS-PAGE gels of Figs 1C and 3C and 4D and S6B and S8A.** (PDF). Contains raw images for Fig 1C, Fig 3C, Fig 4D, S6B Fig, and S8A Fig.

## Notes

https://www.rcsb.org/structure/7ULW

### Competing Interest Statement

The authors have declared no competing interest.

### Summary of Updates

The title, main figures, and supplemental files are updates.

